# Wheat plant height locus *RHT25* encodes a PLATZ transcription factor that interacts with DELLA (RHT1)

**DOI:** 10.1101/2023.01.05.522836

**Authors:** Junli Zhang, Chengxia Li, Wenjun Zhang, Xiaoqing Zhang, Youngjun Mo, Gabriela E. Tranquilli, Leonardo S. Vanzetti, Jorge Dubcovsky

## Abstract

Plant height is an important agronomic trait with a significant impact on grain yield, as demonstrated by the positive effect of the *REDUCED HEIGHT* (*RHT*) dwarfing alleles (*Rht1b*) on lodging and harvest index in the “Green Revolution” wheat varieties. However, these gibberellic acid (GA) insensitive alleles also reduce coleoptile length, biomass production, and yield potential in some environments, triggering the search for alternative GA-sensitive dwarfing genes. Here we report the identification, validation and characterization of the gene underlying the GA-sensitive dwarfing locus *RHT25* in wheat. This gene, designated as *PLATZ-A1 (TraesCS6A02G156600*), is expressed mainly in the elongating stem and developing spike and encodes a plant-specific AT-rich sequence- and zinc-binding protein (PLATZ). Natural and induced loss-of-function mutations in *PLATZ-A1* reduce plant height and its over-expression increases it, demonstrating that *PLATZ-A1* is the causative gene of *RHT25*. PLATZ-A1 interacts physically and genetically with RHT1 (DELLA), and both genes have stronger effects on plant height in the presence of the wildtype than in the presence of the mutant allele of the other gene. These results suggest that PLATZ1 can modulate the effect of DELLA on wheat plant height. We identified four natural truncation mutations and one promoter insertion in *PLATZ-A1* that are more frequent in modern varieties than in landraces, suggesting positive selection during wheat breeding. These mutations can be used to fine-tune wheat plant height and, in combination with other GA-sensitive dwarfing genes, to replace the GA-insensitive *Rht1b* alleles to search for grain yield improvements beyond those of the Green Revolution varieties.

**Significance Statement:** We have identified and characterized a previously unknown gene controlling plant height in wheat and named it *PLATZ1*. Mutations in *PLATZ1* reduce plant height while its overexpression results in taller plants. *PLATZ1* is expressed mainly in elongating stems and developing spikes and interacts physically and genetically with the “Green Revolution” dwarfing gene *REDUCED HEIGHT 1* (*RHT1*). We discovered five natural mutants in the A genome copy of *PLATZ1* in common wheat that have been favored during breeding, suggesting an overall positive effect on wheat performance. These mutations can be used to fine-tune wheat plant height and, eventually, to replace the *RHT1* dwarfing alleles that impose limitations on planting depth and grain yield potential in some environments.

## Introduction

Wheat is a critical crop for global food security and continuous increases in grain yield are required to feed a growing human population that recently surpassed 8 billion people. The large increases in wheat grain yield obtained by the introduction of the *REDUCED HEIGHT* (*RHT*) dwarfing alleles *Rht-B1b* and *Rht-D1b* during the “Green Revolution” demonstrated that optimizing plant height is critical to reduce lodging and improve harvest index (1). *RHT1* encodes a protein designated DELLA (Asp-Glu-Leu-Leu-Ala), which is a critical component of the gibberellin (GA) growth-stimulating pathway (2). GA binds the GID1 (GIBBERELLIN INSENSITIVE DWARF1) receptor altering its conformation and favoring its interaction with the N-terminal region of DELLA, which promotes its subsequent degradation, the release of DELLA-repressed targets, and GA-mediated growth responses (3, 4). Although both *Rht-B1b* and *Rht-D1b* alleles carry premature stop codons in the N-terminal region, DELLA proteins translation is reinitiated after the GID1-GA binding region resulting in GA-insensitive constitutively active repressors (5).

Wheat plants that carry the *Rht-B1b* and *Rht-D1b* alleles accumulate the growth-repressing DELLA proteins, which reduces the stimulating effect of the GA hormone on plant growth (6). As a result, *RHT1b* semidwarf varieties have shorter coleoptiles and reduced above-ground biomass (7). The shorter coleoptiles limit planting depth and access to soil moisture, whereas the reduced biomass can be detrimental in water-limited environments (8). Reduced biomass can also limit grain yield potential in optimum environments, particularly in varieties with excess ‘sink’ capacity (e.g. grain number and size) where a limited ‘source’ (e.g. biomass) can become the limiting factor. Significant progress has been made in recent years in the identification of new alleles for increased grain number (9–11) and grain size in wheat (12), so further increases in biomass are required to fill this extra ‘sink’ capacity. This has triggered new research interests in identifying and testing new GA-sensitive *RHT* genes to replace the GA-insensitive *RHT1* alleles.

The four GA-sensitive wheat *RHT* genes cloned and validated so far show a diversity of gene classes and mechanisms. *RHT8 (TraesCSU02G024900*) encodes an RNase H-like protein carrying a zinc finger BED-type motif (RNHL-D1), and the truncated protein encoded by the *rht-D8* mutant allele is associated with reduced plant height (13, 14). The semidwarf allele *Rht-B13b* (MSTRG.55039) encodes an autoactive nucleotide-binding site / leucine-rich repeat (NB-LRR) protein that causes reduced height due to transcriptional upregulation of pathogenesis-related genes (15). The other two *RHT* genes affect the expression of GA2-oxidase genes that regulate plant height by inactivating endogenous bioactive GA isoforms. The *RHT12* allele for reduced plant height is associated with increased expression of *GA2ox-A14* (16, 17), whereas the related *RHT14, RHT18* and *RHT24* loci are all associated with higher expression of *GA2ox-A9* and reduction of bioactive GA levels in the stems (18, 19).

We previously mapped *RHT25 (QHt.ucw-6AS*) within a 0.2 cM interval on the short arm of chromosome 6A using a population derived from the cross between common wheat lines UC1110 and PI 610750 (20). This genetic interval defined a 4.3 Mb region (144.0 to 148.3 Mb) including 26 high-confidence annotated genes in the Chinese Spring (CS) RefSeq v1.1, but the causative gene was not identified (20). The *RHT25* candidate gene region does not overlap with any of the previously named *RHT* genes on chromosome 6A (20). In this study, we report the identification and validation of the *RHT25* causative gene and its physical and genetic interaction with DELLA. We also describe *RHT25* natural variation and the potential use of these natural alleles to fine-tune plant height in wheat improvement.

## Results

### Identification of *RHT25* candidate gene

To identify the most likely candidate gene for the *RHT25* locus, we took advantage of several mapping populations whose parental lines were previously genotyped by exome capture (21) and characterized for plant height. These included the UC1110 x PI 610750 population used for the initial mapping of *RHT25* (20) and eight recombinant inbred line (RIL) populations generated by crossing the variety ‘Berkut’ to eight different parents (Table 1). These eight populations are part of a nested association mapping (NAM) population previously evaluated for plant height in multiple environments (22, 23).

**Table 1.** Polymorphisms among parents of the NAM populations and the original population used to map *RHT25* (UC1110 x PI 610750). Only polymorphisms within the coding regions and splice sites of genes located in the *RHT25* candidate gene region are presented here. For the *Rht25* alleles, Tall = no significant differences in plant height with PI 610750 or Berkut, and short = significant differences with the previous two parents. A complete list of the SNPs and the haplotypes in this region is available in *SI Appendix*, Table S1. The prioritized candidate gene *TraesCS6A02G156600* is indicated in bold.

| Gene ID | SNP 6AS | Effect | UC1110<sup>1</sup> | PI 610750<br>Berkut | RAC875 | UC1036<br>Dharwar Dry<br>PI 70613 | Cltr 7635<sup>1</sup><br>PBW343<br>RSI5<sup>1</sup> | P515HP<sup>2</sup> |
| --- | --- | --- | --- | --- | --- | --- | --- | --- |
| RefSeq v1.1 | Coordinate | *Rht25* allele: | Short | Tall | Tall | Tall | Tall | Short |
| *TraesCS6A* |  | Haplotype: | H1 | H2 | H3 | H4 | H5 | H6 |
| 02G155900 | 144717100 | L215L | G | G | A | G | G | G |
| **02G156600** | 145651904/16 | Frame shift | 13del<sup>3</sup> | WT | WT | WT | WT | WT |
| **02G156600** | 145651971 | Splice site | T | T | T | T | T | <u>C</u> |
| 02G156700 | 145739155 | R12C | A | G | G | G | G | G |
| 02G157200 | 146501774 | A44A | C | C | C | C | C | <u>A</u> |
| 02G157500 | 147305219 | W165R | A | A | A | A | A | <u>G</u> |

<sup>1</sup> RSI5 and Cltr 7635 have a promoter insertion.
<sup>2</sup> The SNPs specific for P515HP are underlined.
<sup>3</sup> The 13-bp deletion was tested in all accessions with molecular markers described in *SI Appendix*, Table S2.

In the original population used to map *RHT25*, UC1110 was associated with the dwarfing allele and PI 610750 with the allele for increased plant height that is partially dominant over the dwarfing allele (20). Among the eight NAM populations, we only detected significant effects for plant height associated with the *RHT25* locus in the Berkut x Patwin-515HP (henceforth P515HP) population, where P515HP carries the allele for reduced plant height and Berkut the allele for increased plant height. The absence of significant differences in plant height associated with the *RHT25* region in the other seven NAM populations suggests that their parental lines carry the same allele as Berkut (Table 1). Using exome capture data for the parental lines (https://triticeaetoolbox.org/wheat/), we identified 17 SNPs and 2 indels in the *RHT25* candidate gene region that defined six haplotypes (*SI Appendix*, Table S1). Within this region, *TraesCS6A02G156600* was the only gene that showed loss-of-function polymorphisms in the two haplotypes (H1 and H6) associated with the *RHT25* dwarfing allele (Table 1).

In UC1110 (H1), *TraesCS6A02G156600* shows a 13-bp deletion in the third exon (CS 6A: 145,651,904 - 145,651,916) that generates a shift in the reading frame and eliminates 59 % of the predicted amino acids. We developed a marker for this deletion (PlzAF2/PlzAR1, *SI Appendix*, Table S2) and confirmed its presence in UC1110 and its absence in other accessions in Table 1. P515HP (H6) has a mutation in the acceptor splice site of *TraesCS6A02G156600* second intron (CS 6A 145,651,971), which results in the retention of the second intron and a premature stop codon that eliminates 66 % of the predicted amino acids. The other nine accessions carrying the *RHT25* allele for increased plant height showed no polymorphisms in the coding region of *TraesCS6A02G156600*. Moreover, no other gene in the *RHT25* candidate region showed polymorphisms differentiating the two accessions associated with the *RHT25* dwarfing allele from the other accessions carrying the *RHT25* allele associated with tall plants (Table 1). Based on these results, we concluded that *TraesCS6A02G156600* is the most likely candidate gene for *RHT25*, and prioritized it for functional characterization and validation.

### *PLATZ* phylogeny and nomenclature

*TraesCS6A02G156600* encodes a protein previously designated as ‘plant-specific AT-rich sequence- and zinc-binding protein’ (*PLATZ*) (24). A previous phylogenetic study of the 62 *PLATZ* genes identified in wheat (*TaPLATZ1* to *TaPLATZ62*) revealed six groups (I to VI) (25), with *TraesCS6A02G156600 (TaPLATZ34*) included in Group III (25). The two previously characterized wheat *PLATZ* genes belong to Group I (*TaPLATZ5, TraesCS2D02G447400*) (26) and Group II (*TaFl3, TraesCS3A02G497900*) (27), so no functional information is currently available for wheat genes from Group III.

Unfortunately, the gene numbering used in the previous phylogenetic study does not comply with the rules of wheat nomenclature, which require the designation of homoeologous genes with the same identification number (28). Therefore, we performed a new phylogenetic analysis for the wheat PLATZ proteins from Group III (*SI Appendix*, Figs. S1 and S2), identified homoeologs, and assigned them new identification numbers following the wheat nomenclature guidelines (*SI Appendix*, Table S3). We also included the closest *Brachypodium distachyon* (*BRADI*), rice (*Os*) and maize (*Zm*) orthologs to facilitate cross-reference to other grass species, and well-studied homologs from *Pisum* (*Ps*) (24) and *Arabidopsis* (*At*) (29) to facilitate functional comparisons.

Our phylogenetic analysis of the Group III PLATZ proteins showed four well-defined clades designated hereafter as PLATZ1 to PLATZ4, with *TraesCS6A02G156600* included in the *PLATZ1* homoeologous group (*SI Appendix*, Figs. S1 and S2 and Table S3). Orthologs from the four grass-species were present in each of the four clusters, suggesting that they differentiated before the grass radiation. PLATZ1 and PLATZ2 are closely related to each other and to the well-studied PsPLATZ1 (24) and AtORESARA15 (29) proteins, whereas PLATZ3 and PLATZ4 are more divergent.

### Validation of *PLATZ-A1* function using ethyl methanesulfonate (EMS) mutants

To confirm the effect of *PLATZ1* on wheat plant height, we characterized loss-of-function mutations in both *PLATZ-A1* and *PLATZ-B1 (TraesCS6B02G282400LC*) found in the sequenced Kronos EMS population (21). For *PLATZ-A1*, we identified mutant K2557 that carries a mutation in the acceptor splice site of the 3^rd^ intron (RefSeqv1.1 chromosome 6A: 145,651,705). For *PLATZ-B1*, we identified mutant K86 that carries a mutation predicted to change an arginine at position 69 to cysteine (R69C) in the second exon (RefSeqv1.1 chromosome 6A: 207,805,286, *SI Appendix*, Fig. S3*A*). The R69C mutation is located within the PLATZ conserved domain (pfam 04640) and is expected to have a very disruptive effect in protein structure and function based on a negative BLOSUM62 (−3) and very low SIFT score (0) (*SI Appendix*, Method S3).

To test the effect of the splice site mutation in *PLATZ-A1*, we sequenced its transcripts from mutant K2557 using primers Platz6A-F2 and Platz6A-R1 (*SI Appendix*, Table S2) and found two different transcripts. The first and more abundant variant (*SI Appendix*, Fig. S3*B*) showed retention of the third intron, which generates a premature stop codon and the loss of 46.5% of the predicted amino acids. The less abundant variant uses the alternative 2^nd^ next AG splicing site within exon 4, which results in the elimination of 21 bp and the deletion of amino acids VDHVVEQ between positions 131 to 137. The deleted amino acids are located in a very conserved region of the PLATZ1 and PLATZ2 proteins, likely affecting protein function (*SI Appendix*, Fig. S2).

We crossed K2557 and K86 to non-mutagenized Kronos plants five times to reduce background mutations and intercrossed them to generate a BC_4_F_2_ population segregating for mutations in both homoeologs. Compared with the wildtype, the double mutant (henceforth *platz1*) showed a significant (*P* < 0.0001) reduction in plant height (12.7%), peduncle length (18.7%), and other internodes (Fig. 1A and *B* and *SI Appendix*, Table S4). A factorial ANOVA using the two homoeologs as factors showed significant effects in plant height and peduncle length (*P* < 0.0001) for *PLATZ-A1*, but marginally non-significant effects for *PLATZ-B1* (*SI Appendix*, Table S4).

**Figure 1.**
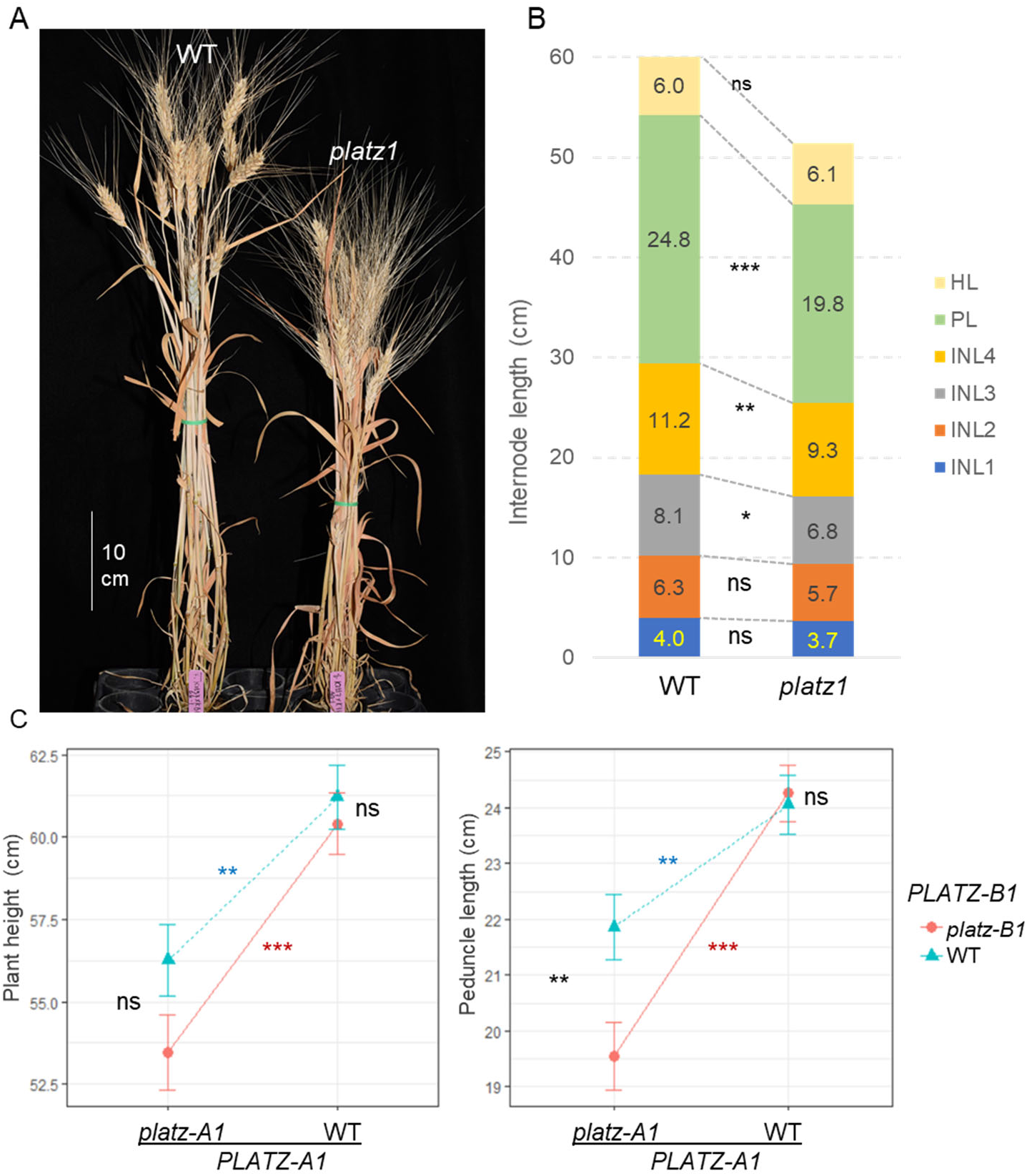
Effect of *PLATZ1* induced mutations on plant height and peduncle length. (**A**) Plant height difference in representative plants. WT= wildtype and *platz1* = double mutant *platz-A1 platz-B1*. (**B**) Differences between wildtype and *platz1* in head length (HL), peduncle length (PL) and internodes length (INL). (**C**) Interaction graphs for *PLATZ-A1* and *PLATZ-B1* (non-parallel lines reflect interactions). Error bars are s.e.m. based on n = 7 to 17. ns= not significant, ** = *P* < 0.01, and *** = *P* < 0.001 for simple effect contrasts in the factorial ANOVA (*SI Appendix*, Table S4).

Since the interaction for peduncle length was significant (*P =* 0.0279, *SI Appendix* Table S4), we analyzed the effects of each gene in the wildtype and mutant backgrounds of the other homoeolog using statistical contrasts (Fig. 1*C*). *PLATZ-A1* showed significant effects on peduncle length under both *PLATZ-B1* backgrounds, but the effects were stronger in the presence of *platz-B1*. Similarly, the *platz-B1* allele showed stronger effects on peduncle length in the presence of *platz-A1* mutant allele (Fig. 1C). These results indicate that *PLATZ-A1* and *PLATZ-B1* have redundant roles in the regulation of plant height and peduncle length, with a stronger effect of *platz-A1* than *platz-B1*.

### Null mutants using CRISPR-Cas9

To test if the smaller effect of the *platz-B1* mutant was the result of a reduced effect of the single amino acid change, we generated truncation mutants in both *PLATZ1* homoeologs using CRISPR-Cas9. We identified a transgenic plant carrying a 1 bp deletion in *PLATZ-A1* that causes a reading frame shift and the truncation of 64% of the protein (*CRplatz-A1*). In the same plant, we identified a 1 bp insertion in *PLATZ-B1* that causes a reading frame shift and the loss of 63% of the protein (*CRplatz-B1*, *SI Appendix*, Fig. S3A). We crossed this transgenic plant with wildtype Kronos and selected plants without the CRISPR-Cas9 vector and carrying both CRISPR-induced mutations.

The F_3_ plants homozygous for the wildtype, *CRplatz-A1, CRplatz-B1* and combined *CRplatz1* mutant showed similar differences in plant height and peduncle length to those described for the EMS mutants in Fig. 1C (*SI Appendix*, Fig. S4 and Table S5). In the presence of *CRplatz-A1*, the *CRplatz-B1* mutation showed a slightly larger effect on plant height (5.1 cm) and peduncle length (3.4 cm) than the *platz-B1* R69C mutation (plant height = 2.8 cm and peduncle length = 2.3 cm), suggesting that the *platz-B1* EMS mutant may have some residual function.

In summary, the combined loss-of-function of the two *PLATZ1* homoeologs resulted in significant reductions in plant height and peduncle length (Fig. S4), indicating that this gene is required to maintain the wildtype plant height.

### Transgenic plants over-expressing *PLATZ-A1* with an HA tag

To test if *PLATZ1* was sufficient to induce changes in plant height, we generated five independent transgenic lines in Kronos overexpressing a fusion of the coding sequence of *PLATZ-A1* and a C-terminal 3xHA tag under the maize *UBIQUITIN* promoter (henceforth, *UBI::PLATZ1-HA*). In leaves of 16-day-old T1 transgenic plants, the combined transcript levels of *PLATZ1* transgene and endogenous genes (3 to 16-fold *ACTIN*) were significantly higher than in the non-transgenic sister lines (0.1-fold *ACTIN, SI Appendix*, Fig. S5). We also observed significant differences in *PLATZ1* expression in peduncle samples collected close to heading time from selected T2 sister lines with and without the transgene from four independent events (*P =* 0.002). In this tissue, transcript levels of *PLATZ1* varied from 43 to 67-fold of *ACTIN* in the presence of the transgene, but were low in the non-transgenic sister lines (0.05 to 0.09-fold *ACTIN, SI Appendix*, Table S6). In spite of the high transcript levels of *PLATZ1* in the *UBI::PLATZ1-HA* transgenic plants, we failed to detect significant differences in plant height in the T2 progenies (*SI Appendix*, Table S7A).

To test if the limited effect of the *UBI::PLATZ1-HA* transgene on plant height was due to sufficient levels of the endogenous gene, we crossed one Kronos *UBI::PLATZ1-HA* transgenic plant with the *platz1* combined mutant and evaluated the F_3_ progeny in a greenhouse experiment. In the absence of the transgene, plants homozygous for the *platz1* mutation were significantly shorter than the non-transgenic wildtype, indicating that the effects of the mutation were detectable under the greenhouse conditions. In contrast, the plants homozygous for the *platz1* mutation carrying the *UBI::PLATZ1-HA* showed an intermediate height and peduncle length and no significant differences with the wildtype (Fig. 2A-B), indicating partial complementation. Compared to the *platz1* mutant, the plants that carry the mutation and the transgene showed a significantly longer peduncle (*P* = 0.018); yet the increase in plant height (5.25 cm) was not significant (Fig. 2B, *SI Appendix*, Table S7B). In summary, the *UBI::PLATZ1-HA* transgene was able to partially complement the effect of the *platz1* mutant on both peduncle length and plant height.

**Figure 2.**
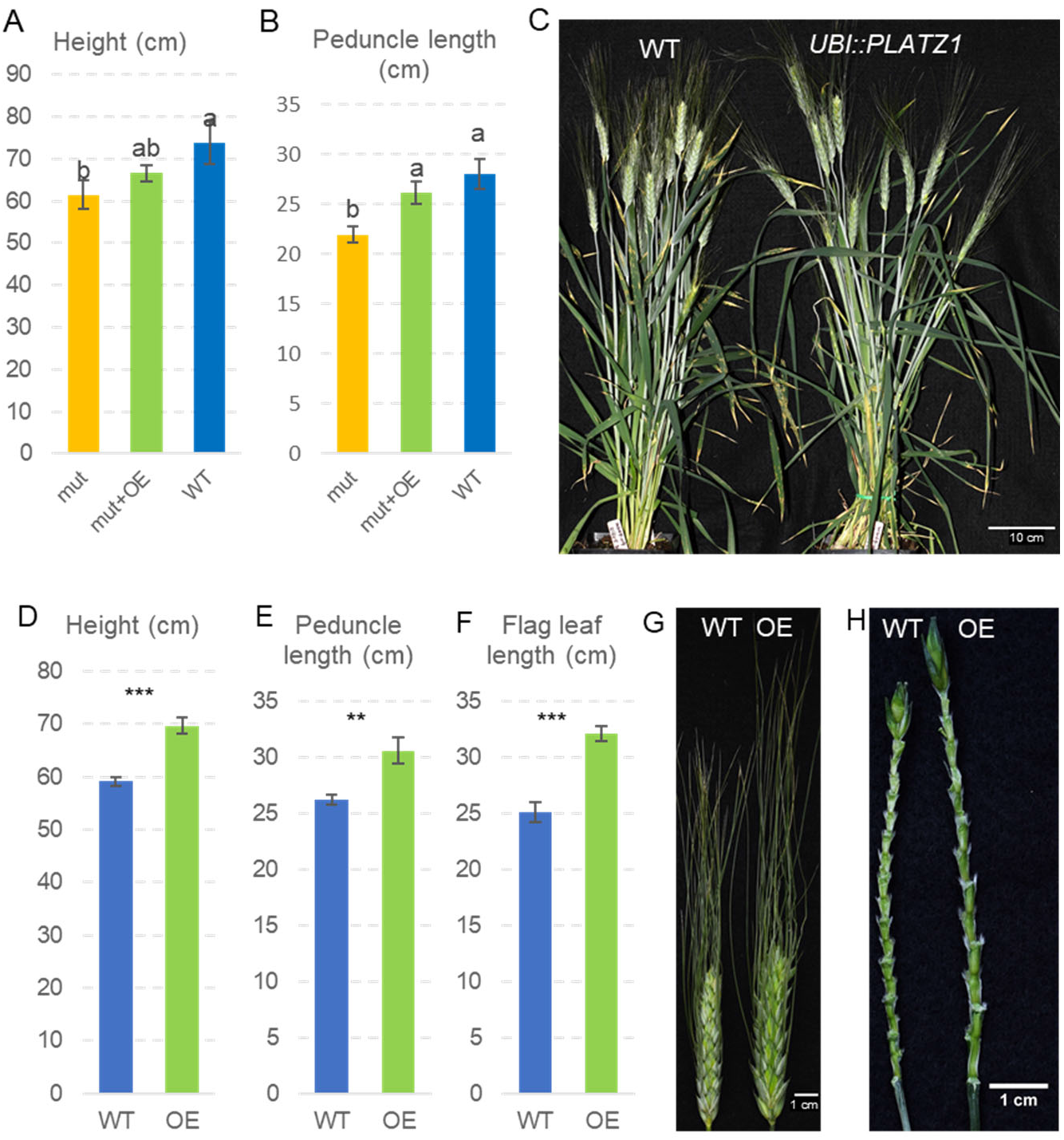
Effect of *UBI::PLATZ1-HA* and *UBI::PLATZ1* transgenes on plant height and peduncle length. (**A-B**) Complementation experiment in sister lines carrying the wildtype allele (WT), the *platz1* double mutant (mut), and *platz1* combined with the *UBI::PLATZ1-HA* transgene (mut + OE). Error bars are s.e.m. and different letters above the bars indicate significant differences (*P* = 0.05, n = 6, *SI Appendix*, Table S7B). (**C-H**) Kronos BC_1_F_1_ plants transformed with the construct *UBI::PLATZ1* without the HA tag (OE n = 8 and WT n = 5, *SI Appendix*, Table S7C). (**C**) Selected plants showing differences in plant height. (**D**) Plant height. (**E**) Peduncle length. (**F**) Flag leaf length. ** = *P* < 0.01 and *** = *P* < 0.001. (**G**) Spikes showing fewer but larger spikelets in the *UBI::PLATZ1* transgenic plants (OE) than in the wildtype (WT). (**H**) Spikes without the spikelets showing fewer but longer internodes in the plants carrying the *UBI::PLATZ1* transgene (OE) than in the wildtype (WT). These transgenic plants were mostly sterile.

Taken together, the complementation and mutant experiments demonstrate that *PLATZ-A1* is the causative gene of *RHT25*.

### Transgenic plants over-expressing *PLATZ-A1* without a tag

To test if the presence of the HA tag was altering the effect of the transgene, we generated additional transgenic plants with a similar *UBI::PLATZ1* construct but without the tag. The T0 plants were mostly sterile, so we crossed them with wildtype Kronos and analyzed the phenotype in F_1_ and BC_1_F_1_ plants with and without the transgene (Fig. 2C-H, *SI Appendix*, Table S7C).

The F_1_ and BC_1_F_1_ plants carrying the transgene showed taller plants (Fig. 2C-D), longer peduncles (Fig. 2E), and longer flag leaves (Fig. 2F). Some of the lower leaves were also longer and showed reduced width (*SI Appendix*, Table S7C). In addition, the transgenic plants headed later, had spikes with fewer spikelets and longer rachilla internodes (Fig. 2G-H). The spikelets were larger (Fig. 2G) as reflected by significantly longer glumes (*SI Appendix*, Table S7C).

In summary, the transgenic plants showed more drastic phenotypes than the plants transformed with the *UBI::PLATZ1:HA*, suggesting that the HA tag partially interfered with the PLATZ1 function.

### *PLATZ1* expression

Using previously published RNA-seq data, we found that *PLATZ1* transcript levels in hexaploid wheat were the highest in the stem during the early stages of elongation (Z30-Z32, Zadoks’ scale) followed by the early developing spikes (Z32). Transcripts of *PLATZ1* were also observed in the leaves, roots and grains, but at lower levels (30). In most tissues, *PLATZ1* transcript levels were the highest in the D-genome homoeolog, intermediate in the A-genome homoeolog, and the lowest in the B-genome homoeolog (*SI Appendix*, Fig. S6A). We also observed a relative higher abundance of *PLATZ-A1* compared to *PLATZ-B1* transcripts at different stages of spike development in RNA-seq data for tetraploid wheat Kronos (Fig. S6B) (31). We also explored *PLATZ-A1* transcript levels in different sections of the elongating peduncle in Kronos plants at ear emergence (Zadok 55) using qRT-PCR. The transcript levels of *PLATZ-A1* in sections including the node and 1 cm immediately above the node were significantly higher (Tukey test, *P* < 0.05) than in sections starting 2 cm above the node (*SI Appendix*, Table S8 and Fig. S6C).

We then explored the accumulation profile of the protein encoded by the *UBI::PLATZ1-HA* transgene. First, we performed transient expression in Kronos protoplasts and detected abundant levels of PLATZ1-HA protein in western blots using an anti-HA antibody (Fig. 3A, lane 9). However, when we extracted total proteins from the 2^nd^ fully expanded leaf of Kronos T3 transgenic plants at the 3-leaf stage, we were unable to detect the protein encoded by the *UBI::PLATZ1-HA* transgene (Fig. 3A, lanes 1-2), despite high transcript levels (Fig. S5 and Table S6).

**Figure 3.**
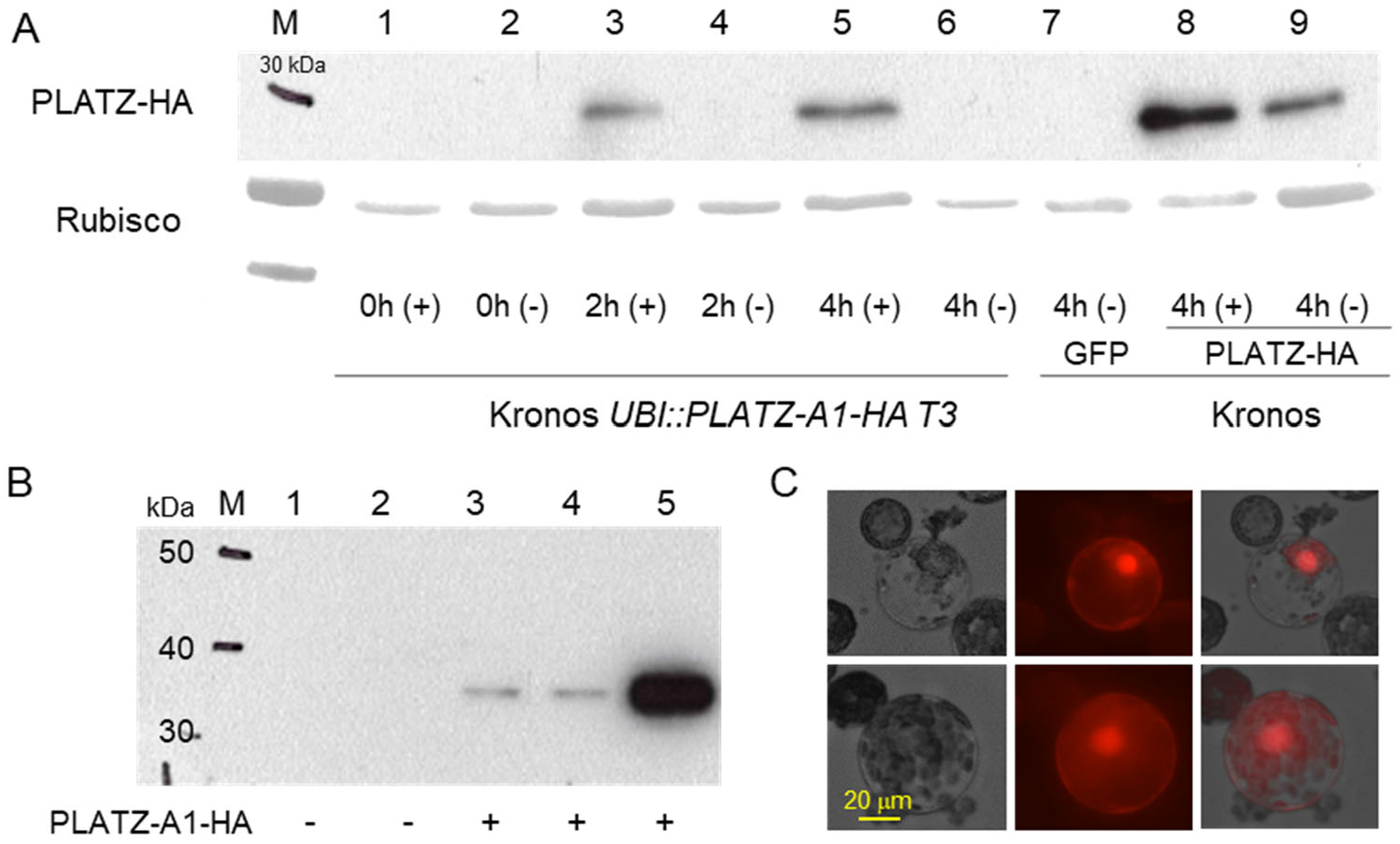
Western blots of PLATZ1-HA using an anti-HA antibody. **A**) Lanes 1-6 are protoplast prepared from T3 transgenic *UBI::PLATZ1-HA* plants treated with either 50 μM MG132 solution in DMSO (+) or only DMSO (−). Protein was extracted 0 h, 2 h and 4 h after treatment. Lanes 7-9 are Kronos protoplasts transiently transformed with *UBI::GFP* (7) or *UBI::PLATZ1:HA* (8–9) with (8) or without MG132 (9). Protein was extracted 4 h after treatment. **B**) Western blots of proteins extracted from the shoot apical meristem (SAM) and stems. Lanes 1 and 2 are from non-transgenic Kronos plants (negative control), lanes 3-4 are from a Kronos *UBI::PLATZ1-HA* T3 plant and lane 5 is from transiently transformed Kronos protoplasts (positive control). Lanes 1 and 3 include the SAM and the nodes below collected before elongation (4^th^ leaf stage, Waddington scale 3.0). Lanes 2 and 4 are from stems collected at the early elongation stage. The expected size of the PLATZ1-HA protein is 32 kDa. **C**) Nuclear localization of an mCherry-PLATZ-A1 chimera.

To test if protein degradation was responsible for our inability to detect the PLATZ1-HA protein, we treated protoplasts extracted from leaves of the transgenic *UBI::PLATZ1-HA* plants with the proteasome-inhibitor MG132 solution in DMSO or only DMSO as a negative control. Two hours after the treatment, we detected the PLATZ1-HA protein in the protoplasts treated with MG132 but not in those treated with DMSO, and also the signal became stronger four hours after the treatment (Fig. 3A). These results suggest that the PLATZ1-HA protein is rapidly degraded in the leaves through the proteasome pathway. In control leaf protoplasts from the non-transgenic Kronos plants transiently transformed with *UBI::PLATZ1-HA*, protein levels were higher in the presence of MG132 than in its absence (Fig. 3A, lanes 8 and 9), suggesting that the protein was also degraded by the proteasome in the protoplasts. Our western blot analyses detected much higher levels of PLATZ1-HA protein in the transiently transformed protoplasts than in the leaves (both in the presence and absence of MG132), which is likely due to the high concentration of plasmid DNA used for protoplast transfection.

Since the endogenous *PLATZ1* gene is expressed at higher levels in the elongating stems than in the leaves (Fig. S6A), we tested if we could detect the PLATZ1-HA protein in the shoot apical meristem (SAM) and stems of the *UBI::PLATZ1-HA* transgenic plants in the absence of MG132. In samples extracted from the SAM and the stem nodes before elongation and in a later stage from early elongating nodes, we detected PLATZ1-HA in the *UBI::PLATZ1-HA* transgenic plants but not in the non-transgenic negative controls (Fig. 3B).

Finally, we expressed a *p2xCa35S::mCherry-PLATZ* chimera in the Kronos wheat protoplast to determine its subcellular localization. The PLATZ1 proteins were mainly located in the nucleus (Fig. 3C), an expected subcellular localization for a transcription factor.

### Genetic and physical interactions between PLATZ1 and DELLA

To test if *PLATZ-A1 (RHT25*) genetically interacts with *RHT1*, we crossed the *platz-A1* mutant with a tall Kronos near-isogenic line in which the *Rht-B1b* dwarfing allele was replaced by the *Rht-B1a* allele. We selected plants homozygous for the four possible allele combinations in the F_2_ progeny and evaluated the F_3_ progeny for plant height and peduncle length (*SI Appendix*, Table S9). A factorial ANOVA showed significant effects for both *PLATZ-A1* and *RHT1* on plant height and peduncle length, and significant interactions for plant height (*P =* 0.011) and marginally non-significant for peduncle length (*P =*0.061, *SI Appendix*, Table S9).

We then tested if the significant genetic interaction between *RHT25* and *RHT1 (DELLA*) was associated with a physical interaction between their encoded proteins using yeast-two-hybrid (Y2H) assays. We used a truncated DELLA protein carrying only the C-terminal GRAS domain (RHT1-GRAS) because the fulllength protein showed autoactivation in Y2H assays. PLATZ-A1 showed a positive interaction with RHT1-GRAS (Fig. 4A). We further dissected the RHT1-GRAS domain into three sub-domains and evaluated their interactions with PLATZ1. We observed a positive interaction between PLATZ1 and the PFYRE subdomain, but no interaction with the LVL and SAW sub-domains in the Y2H assays (Fig 4B).

**Figure 4.**
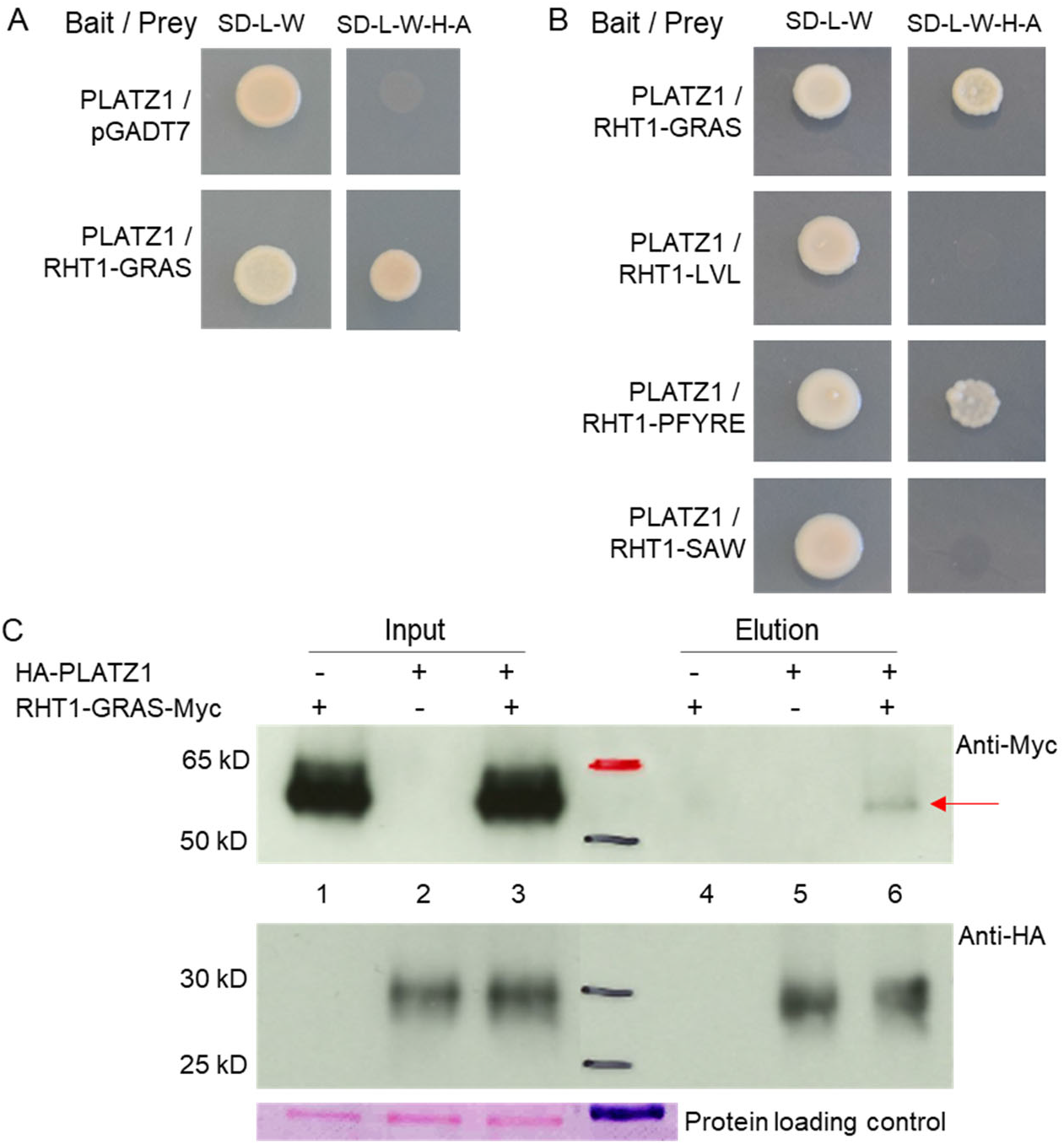
Interactions between PLATZ-A1 and DELLA (RHT1). **A**) Yeast-two-hybrid (Y2H) assays between DELLA and PLATZ-A1. **B**) Y2H between the complete PLATZ-A1 protein and three subdomains of the DELLA-GRAS protein (LVL, PFYRE, and SAW). **C**) Co-IP assay between PLATZ-A1 and RHT1-GRAS in wheat protoplasts. The pull-down assay was carried out with anti-HA magnetic beads. Ponceau S staining was used as loading control. The red arrow indicates the RHT1-GRAS-Myc protein coprecipitated with HA-PLATZ1.

A co-immunoprecipitation (Co-IP) experiment using Kronos wheat protoplasts validated the interaction between PLATZ-A1 and DELLA-GRAS *in vivo*. We co-transformed HA-PLATZ1 and RHT1-GRAS-Myc into Kronos protoplasts and, after immunoprecipitation with anti-HA beads, we detected a weak signal for RHT1-GRAS-Myc with the anti-Myc antibody (Fig. 4C).

DELLA has been reported to physically interact with the GROWTH-REGULATING FACTOR 4 (GRF4) (6), so we evaluated the 3-way interactions between DELLA, PLATZ1 and GRF4 using yeast-three-hybrid (Y3H) assays. We first determined that GRF4 does not exhibit autoactivation, and that it can interact with DELLA but not with PLATZ-A1 in Y2H assays (Fig. 5A). We found that expression of GRF4 as the 3^rd^ protein in quantitative alpha-gal Y3H assays, reduced the interaction between PLATZ1 and DELLA by 74% (Fig. 5B, *P* < 0.001). In contrast, when PLATZ-A1 was expressed as the 3^rd^ protein in the Y3H assay, it did not produce a significant effect (*P* = 0.58, *SI Appendix*, Table S10) on the interaction between GRF4 and DELLA (Fig. 5C). This result is consistent with the >20-fold stronger interaction between GRF4 and DELLA than between PLATZ and DELLA (Fig. 5B-C). Taken together, these results suggest that GRF4 can be a competitor for the DELLA – PLATZ-A1 interaction.

**Figure 5.**
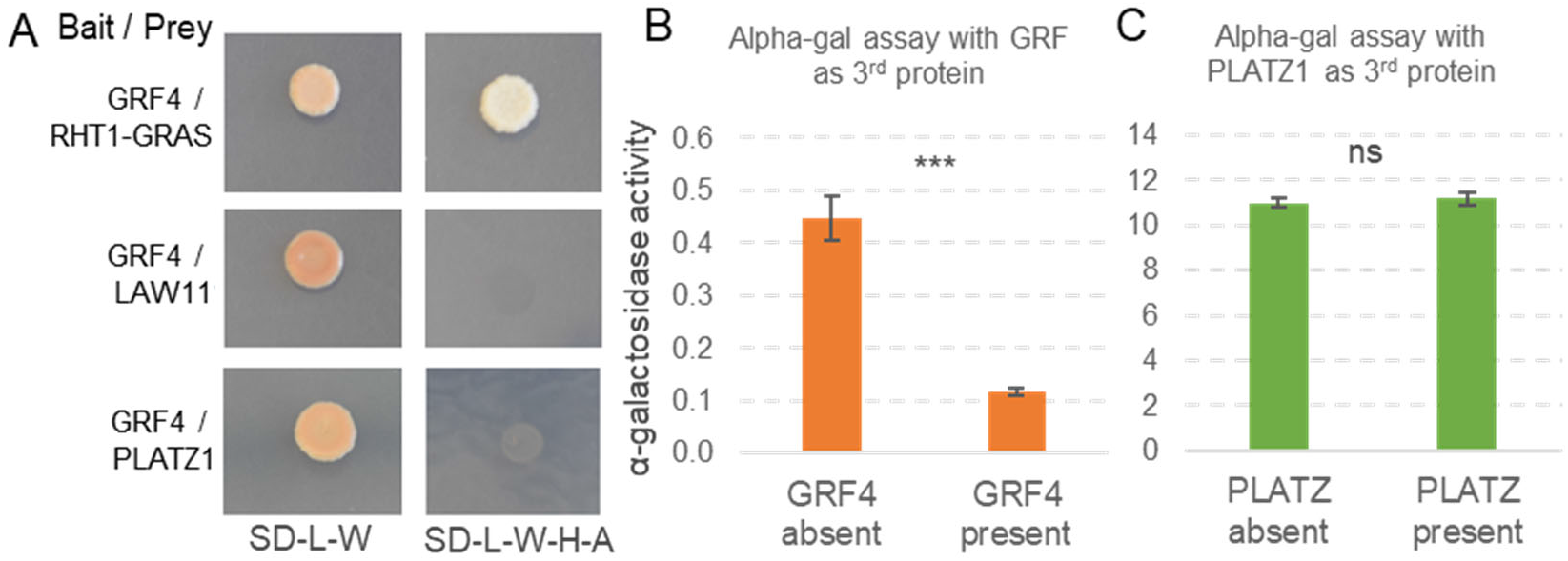
Yeast-three-hybrid (Y3H) assays between GRF4, DELLA, and PLATZ-A1. **A**) Interaction between GRF4 and RHT1-GRAS, PLATZ-A1 and empty vector LAW11 (autoactivation test). **B**) Quantitative alpha-gal Y3H assay between PLATZ-A1(DB) and RHT1 (AD) with GRF4 expressed or not expressed as the 3^rd^ protein. **C**) Quantitative alpha-gal Y3H assay between GRF4 (DB) and RHT1(AD) with PLATZ-A1 expressed or not expressed as the 3^rd^ protein. *** = *P* < 0.001 and ns = not significant. Data in *SI Appendix*, Table S10.

### Natural variation in *PLATZ-A1*

To explore if *PLATZ-A1* dwarfing alleles were selected during wheat domestication and breeding, we determined the frequency of *PLATZ-A1* natural variants using available exon capture data from 45 common wheat lines deposited in the Wheat T3 database and genomic sequences available through the PanGenome Project (32). In addition to the 13-bp deletion found in UC1110 and the splice site mutations detected in P515HP, we found two additional deletions in the third exon of *PLATZ-A1*. The first one is a 4-bp deletion in the reference genome of CS RefSeq v1.1 (CAGG deletion between 6A:145,651,864 and 145,651,865, in - strand) and the second one is a 19-bp deletion (CS RefSeq v1.1, 6A:145,651,892 - 145,651,910). Finally, we detected a 384-bp insertion in the promoter region 130 bp upstream of the start codon in CDC Landmark (CS RefSeq v1.1, 6A:145,654,767 - 145,654,768). The annotated sequence in CDC Landmark indicates a 232 bp insertion relative to CS, but Sanger sequencing and PCR product sizes from CDC Landmark (*SI Appendix*, Fig. S7) confirmed the presence of a 384-bp insertion. The effect of the different mutations on the encoded PLATZ1 protein, together with a haplotype analysis including the *RHT25* natural alleles, is presented in *SI Appendix*, Table S11.

We designed a single pair of primers (PlzAF2/PlzAR1, Table S2) that can detect all four truncation mutations, as well as a KASP marker that detects the promoter insertion (*SI Appendix*, Fig. S7). We used these primers to screen 46 *T. turgidum* ssp. *dicoccoides*, 78 *T. turgidum* ssp. *dicoccon*, 508 *T. turgidum* ssp. *durum (SI Appendix*, Table S12), and 1,120 common wheats (*SI Appendix*, Table S13). The common wheats included 45 modern spring and winter varieties from the Wheat Coordinated Agricultural Project (WheatCAP), 236 accessions from the North America photoperiod insensitive spring wheat association mapping panel (hereafter referred to as the spring panel), and 839 from the USDA National Small Grains Collections (NSGC, *SI Appendix*, Table S13).

The combined *RHT25* mutations represented 30% of the hexaploid accessions, but were completely absent from the ancestral or modern tetraploid wheat accession (*SI Appendix*, Tables S12-13). The frequency of the combined mutant alleles in hexaploid wheat was significantly higher (χ^2^ tests *P* < 0.001) among the advanced materials (29.7 %) than among landraces (5.0 %). Within the advanced materials, the mutant alleles were more frequent among those released after 1960 (28.1 %) than among those released before 1960 (14.9%, *SI Appendix*, Table S13). These results suggest that the *PLATZ-A1* mutant alleles were subject to selection during wheat improvement.

The most frequent mutation in the hexaploid wheat collection was the 13-bp deletion (*Rht25b*, 15.8%), followed by the promoter insertion (*Rht25f*, 11.0%), whereas the other three truncation mutations (*Rht25c-e*) were detected in < 2% of the accessions (*SI Appendix*, Table S13). Both the *Rht25b* and *Rht25f* alleles showed higher frequencies in advanced materials than among landraces (*P* < 0.001), and in materials released after 1960 than before, but the latter was significant only for *Rht25b*. The other truncation alleles showed no significant differences, but the frequencies were too small to compare (*SI Appendix*, Table S13).

### Evaluation of the effect of natural *RHT25* alleles on plant height

We used biparental populations segregating for *Rht25c, Rht25d, Rht25e*, and *Rht25f* to study their effects on plant height (*SI Appendix*, Table S14). To characterize the effect of *Rht25c* (19-bp deletion) on plant height, we used the previously published population McNeal (*Rht25c*) x Thatcher (*Rht25f*) (33). This population showed a QTL for plant height in the *RHT25* region on 6AS and is fixed for the *Rht24a* allele in 6AL. We genotyped 153 RILs from this population with the *PLATZ-A1* marker and confirmed that *Rht25c* was associated with a highly significant decrease (*P* < 0.001) in plant height relative to *Rht25f* in a factorial ANOVA including the *RHT-D1* alleles (*SI Appendix*, Table S14).

To characterize the effect of *Rht25d* (splice mutation), we used the Berkut (*Rht25a*) × P515HP (*Rht25d*) NAM population. We identified an F_5_ line heterozygous for the *PLATZ-A1* region and homozygous for *Rht24b* and derived a heterogeneous inbred family (HIF). We genotyped 62 F_5:2_ HIF lines with the *PLATZ-A1* marker and found a highly-significant effect on plant height associated with the *Rht25d* allele (8.6 cm, *P* < 0.001, *SI Appendix*, Table S14).

To study the effect of *Rht25e* (4-bp deletion) on plant height, we evaluated a population from the cross between RSI5 _(*Yr5+Yr15+Glu-A1a+GPC-B1*)_ (hereafter RSI5, *Rht25f*) and PI 520033 (*Rht25e*). We identified a single F_4_ plant heterozygous for *RHT25* and fixed for *Rht24a, Rht-B1a*, and *Rht-D1a* and tested its F_5_ progeny for plant height in the greenhouse. The plants homozygous for the *Rht25e* allele were 5.5 cm shorter than those homozygous for the *Rht25f* allele (*P* = 0.009). We also compared BC_4_F_2_ homozygous sister lines from the cross between CS (*Rht25e*, used as recurrent parent) and Spring Hobbit (*Rht25a*). Plants homozygous for the Spring Hobbit allele (*Rht25a*) were 9.6 cm taller than those carrying the *Rht25e* allele (*P* < 0.05).

For *Rht25f* (promoter insertion), we developed one F_3:4_ and two F_4:5_ populations from the cross between CDC Landmark (*Rht25f*) and Berkut (*Rht25a*). An ANOVA combining the three families and using family as block showed a 2.1 cm reduction in plant height in the sister lines carrying the *Rht25f* relative to the wildtype but the difference was marginally not significant (*P =* 0.0613). However, two indirect sources of evidence suggest that this small difference is likely real. First, the differences in plant height between *Rht25f* and truncation mutants *Rht25c* (2.8 cm) and *Rht25e* (5.5 cm) were smaller than between the wildtype allele (*Rht25a*) and the truncation mutations *Rht25b* (17.7 cm), *Rht25d* (8.6 cm) and *Rht25e* (9.6 cm, *SI Appendix*, Table S14). In addition, the promoter insertion in *Rht25f* was associated with a 38% decrease in *PLATZ-A1* transcript levels relative to *Rht25a* in the elongating peduncles (*P* < 0.05) in qRT-PCR experiments (*SI Appendix*, Table S15).

### Effect of *PLATZ1* natural and induced mutations on grain size and weight

The rice gene *Os06g0666100*, designated as *GRAIN LENGTH 6* (*GL6*) (34) or *SHORT GRAIN 6 (SG6*) (35) in separate studies, is a close homolog of the wheat *PLATZ1* gene (*SI Appendix*, Fig. S1). Rice plants overexpressing this gene have larger and heavier grains, whereas those carrying mutations in this gene have reduced grain length and weight. Based on these results, we explored the effect of wheat *PLATZ1* on grain size in natural and induced *PLATZ1* mutants (the transgenic wheat plants overexpressing *PLATZ-A1* were mostly sterile).

We first evaluated the effect of *Rht25b* on grain length, grain width and grain area in 186 RILs from the UC1110 x PI 610750 population in field experiments performed in Argentina (25 grains from 5 random spikes for each RIL). The RILs carrying the *Rht25b* allele showed a significant reduction (*P* < 0.01) in grain length (2.0%), grain width (2.6%), grain area (4.5%), and thousand kernel weight (TKW, 6.0%) relative to those carrying the wildtype allele (*SI Appendix*, Table S16A).

We then evaluated the effect of the induced *platz-A1* and *platz-B1* loss-of-function mutations on grain development in tetraploid wheat. We introgressed both mutations into the semidwarf (*Rht-B1b*) commercial durum wheat varieties Kronos, Desert Gold (PVP 2019-00010) and the UC Davis breeding line UC1771-Low Cadmium (UC1771LC) by four backcrosses (BC_4_F_3_), and evaluated them at the University of California, Davis experimental field in 2022 in a randomized complete block design with 5 replications using small plots (4.5 m^2^).

A combined ANOVA including the three lines is presented in *SI Appendix*, Table S16A. As expected, plants carrying the *platz1* mutations showed a significant reduction in plant height relative to the wildtype, with the largest effect in the combined *platz1* mutant (−13.0%), followed by *platz-A1* (−10.5%) and *platz-B1* (−2.5%). The stronger effect of the combined *platz1* mutant on plant height was also reflected in stronger effects on grain length (−1.4%, *P* < 0.001), grain width (−2.9%, *P* < 0.05), grain area (−2.8%, *P* < 0.001), TKW (−4.1%, *P* < 0.01), and grain yield (−5.5%, *P* < 0.05, *SI Appendix*, Table S16B1). The individual mutants showed no significant differences for most traits, except for grain yield in *platz-A1* and grain length in *platz-B1*. In ANOVAs performed separately by variety, the differences in grain yield were not significant and the differences in grain size and weight were significant only in UC1771LC (*SI Appendix*, Table S16B2), suggesting that the *PLATZ1* effects are modulated by the genetic background.

## Discussion

### *PLATZ-A1* is the causative gene for the *RHT25* locus

In this study, we show that among the genes completely linked to *RHT25* (20), *TraesCS6A02G156600* is the only one with polymorphisms that differentiate tall and semidwarf *RHT25* alleles in two different segregating populations. We also show that induced mutations in *TraesCS6A02G156600* result in reduced height, and that transgenic lines overexpressing this gene partially complement the mutant phenotype. Taken together, these results demonstrate that *TraesCS6A02G156600* is the causative gene underlying the wheat plant height locus *RHT25*.

*TraesCS6A02G156600* encodes a plant-specific PLATZ zinc-dependent DNA-binding protein with two noncanonical zinc finger domains conserved in the wheat PLATZ proteins (*SI Appendix*, Fig. S2). Among the six clusters identified in a phylogenetic analysis of the wheat PLATZ proteins (25), TraesCS6A02G156600 was part of Group III. This group includes four wheat paralogs (*PLATZ1* to *PLATZ4*, *SI Appendix*, Fig. S1), as well as previously characterized *PsPLATZ1* from pea (*Pisum sativum*) (24), its related Arabidopsis homolog *AtORESARA15* (29), and a rice gene known as either *GL6* or *SG6* (34, 35). The multiple functions of these genes are compared in the section below with those observed in this study for wheat *PLATZ1*.

### *PLATZ* genes from Group III have multiple pleiotropic effects

The *PsPLATZ1* gene from pea was the first *PLATZ* gene identified in plants, and its two zinc-finger domains were found to be required for both Zn-binding activity and for binding to A/T-rich DNA sequences (24). Its closest Arabidopsis gene, *AtORESARA15*, was shown to be involved in the regulation of both leaf growth and senescence (29). Loss-of-function mutations in this gene resulted in smaller leaves and accelerated senescence under salt stress, whereas a dominant mutant showed multiple pleiotropic effects including enlarged leaf size, extended leaf longevity, increased plant height and root length, and increased seed volume and weight (29).

Similar effects on seed volume and weight were observed in the rice *PLATZ* gene *Os06g0666100* (also known as *GL6* or *SG6*) (34, 35), which is related to the wheat *PLATZ1* gene (*SI Appendix*, Fig. S1). Rice loss-of-function mutants for this gene have reduced grain length (−10 to −13%) and weight (−14 to −21%) but no significant changes in grain width, whereas transgenic plants with higher *Os06g0666100* expression levels showed significantly longer and heavier grains (34, 35). The wheat grains in the combined *platz1* mutant showed a similar trend, but the effects on grain length (−1.4%) and grain weight (−4.1%, *SI Appendix*, Table S16B) were smaller than those observed in rice. The transgenic wheat line overexpressing *UBI::PLATZ1* was male sterile, so we were not able to test the effect of the transgene on grain size; yet we observed a significant increase in glume size (*SI Appendix*, Table S7C), similar to what was observed in rice transgenic plants.

The rice *gl6* mutant showed a significant increase in the number of spikelets and grains per panicle relative to the wildtype (34), a phenotype observed also in the wheat population segregating for *RHT25*. The *Rht25b* dwarfing allele was also associated with a significant increase in spikelet number per spike (*P* < 0.001), but this increase was not translated into a significant increase in grain number per spike (20). The wheat *UBI::PLATZ1* transgenic plants were male-sterile, a phenotype that may have contributed to the dramatic reduction in grain number observed in the transgenic rice plants overexpressing *Os06g0666100* (34, 35). Similar to the rice *gl6* mutant, the wheat *platz1* mutant showed a reduction in plant height and grain yield. Taken together, these results suggest conserved functions of the *PLATZ* genes from Group III in the wheat and rice lineages.

The multiple pleiotropic effects of these *PLATZ* genes parallel their ubiquitous expression profiles in both wheat (*SI Appendix*, Fig. S6) and rice (34, 35). Although transcripts were detected in all tested tissues, expression was the highest in the early stages of inflorescence development in both species. In wheat, *PLATZ1* was also expressed at high levels in the early stages of stem elongation (*SI Appendix*, Fig. S6) and in the elongating peduncle in the nodes and its adjacent region. This result is consistent with the stronger expression of *Sg6* in the nodes of the elongating stems in rice, where cell division contributes new cells to stem elongation (35).

Wheat and rice also showed expression of these related *PLATZ* genes in the grains. In wheat grains, *PLATZ-A1* is the highest in later stages of development (*SI Appendix*, Fig. S6), whereas in rice *Sg6* is highly expressed in the embryos but not in the endosperm. Finally, *in situ* hybridization of *Gl6* in rice showed expression in stamen primordia and mature anthers (34), which may be related to the male sterility we observed in the *UBI::PLATZ1* wheat lines.

### Proposed action mechanisms of PLATZ proteins from Group III

The shorter spikelet hull and grains in the rice *gl6/sg6* mutants were determined by reduced cell numbers and were associated with the downregulation of multiple cell-cycle-related genes that promote cell proliferation (34, 35). In Arabidopsis, *AtORESARA15* was also found to promote early leaf growth by enhancing the rate and duration of cell proliferation activity. A dominant mutant of this gene (*ore15-1D*) showed an upregulation of genes with positive effects on cell proliferation, such as *Cyclin D3;1, GROWTH-REGULATING FACTOR 5* (*GRF5*) and its cofactor *GIF1;* and a downregulation of negative regulators such as miR396, which targets *GRF* transcription factors for degradation. The encoded AtORESARA15 protein was found to directly promote the expression of *GRF1* and *GRF4* (29).

In wheat, we found a similar set of cell cycle-related genes that were significantly downregulated in the elongating peduncle of the CRISPR *platz1* combined mutant relative to the wildtype. These genes include *GRF1, GRF5, GRF9, GRF10, GRF11*, and CYCD4;1 (−17 to −55% reduction, *P* < 0.5 *SI Appendix*, Table S17). We also observed reduced levels of *GRF3* (−43%, *P =* 0.06), *GRF4* (−30%), *GIF1* (−30%) and cyclins CYCD1;1 (−11%) and CYCD3;1 (−16%, *P* = 0.06) in *platz1*, but those differences were not significant (*SI Appendix*, Table S17). Taken together, these results suggest that reduced cell proliferation likely contributed to the reduced wheat plant height of the *platz1* mutant and the increased length of transgenic *UBI::PLATZ1* stems, leaves and glumes. These results also suggest a conserved role of the *PLATZ* genes from Group III in the regulation of cell proliferation.

The physical interaction between wheat PLATZ-A1 and DELLA reported in this study has not been reported before and provides a potential additional mechanism by which PLATZ proteins can affect plant height. The biological relevance of this physical interaction on plant height is supported by the significant genetic interaction between *RHT25* and *RHT1*. This interaction was observed in hexaploid wheat using the natural *Rht25b* allele (20), and confirmed in tetraploid wheat using the EMS-induced *platz-A1* mutant. These results indicate that *PLATZ1* and *DELLA* operate in a connected pathway regulating plant height.

DELLA proteins have a unique N-terminal GA perception region for binding the GA receptor GID1 and a conserved C-terminal GRAS domain required for its repression activity through interactions with multiple regulatory proteins. Mutations in the wheat DELLA N-terminal region in the *Rht-B1b* alleles result in GAinsensitivity and the accumulation of DELLA, which has negative effects on nitrogen-use efficiency and growth (6, 36). Studies in rice have shown that the negative effects associated with the accumulation of DELLA can be counterbalanced by the opposing activity of the *GRF4* transcription factor that physically interacts with DELLA (6). We observed in wheat a similar interaction between DELLA and GRF4 proteins (Fig. 5A), and show that this interaction is able to interfere with the interactions between PLATZ1 and DELLA proteins in Y3H assays (Fig. 5B). These results suggest that PLATZ proteins from Group III may be involved in the significant role played by the GRF-DELLA interaction on the regulation of plant growth (6).

### *RHT25* natural variation and uses in wheat improvement

None of the mutations in *PLATZ-A1* detected in this study was found in the wild or cultivated tetraploid wheat species, which suggests that they likely had originated in hexaploid wheat. The *Rht25b, Rht25c* and *Rht25f* alleles were found in hexaploid landraces suggesting a relatively old origin. The landraces with the *Rht25b* allele were from Iran, Armenia and Greece; those with the *Rht25c* allele from Georgia and Turkey, and those with the *Rht25f* allele from Turkey and Greece. These results suggest that these mutant alleles likely appeared close to the geographical origin of hexaploid wheat. *Rht25d* was found only in more modern materials, which suggests a more recent origin. The *Rht25e* allele present in Chinese Spring, was detected in a limited number of varieties from diverse locations, so it is difficult to infer a potential origin (*SI Appendix*, Table S13).

The significant increases in the proportion of *PLATZ-A1* mutant alleles in advanced materials relative to landraces, and within the advanced materials in post-1960 relative to pre-1960 releases suggest positive selection pressure for the mutant alleles during wheat breeding. The increased frequency of the mutant alleles also indicates that the benefits of the reduced height of the *platz-A1* alleles likely outweigh the small negative effect on grain size detected in this study. Our field experiment was performed in a semidwarf variety (*Rht-B1b*) and in an environment with optimum resources (see high yields in *SI Appendix*, Table S16B1) where the reduced height may have limited grain yield potential. It will be important to study the effects of the *PLATZ-A1* mutant alleles in different environments and genetic backgrounds.

We initially thought that the *platz-A1* loss-of-function alleles could be more beneficial in tall than in semidwarf wheat varieties. However, a comparison of the frequency of the *PLATZ-A1* truncation mutations in post-1960 varieties showed a significantly higher frequency (χ^2^, *P* < 0.001) among semidwarf (42.6 %) than among tall wheat varieties (23.1%, *SI Appendix*, Table S13), suggesting that our hypothesis may not be correct. We also observed a significantly higher (χ^2^, *P* < 0.001) proportion of *platz-A1* mutations among spring varieties developed under spring planting (71.8 %) than among those developed under fall planting (41.7 %, *SI Appendix*, Table S13). We hypothesize that this difference may be related to the longer cycle and higher yield potential of the fall-planted spring varieties. However, these allele frequencies can be affected by multiple historical factors, so these hypotheses need to be validated experimentally.

One of the motivations for identifying the *RHT25* causative gene was the current interest in wheat breeding programs to replace the *RHT1* GA-insensitive alleles by GA-sensitive ones to improve seed planting depth, nitrogen use efficiency and biomass production (6, 8, 36). However, the *platz-A1* mutant alone reduces height by ~10 to 14%, so they will need to be combined with other GA-sensitive dwarfing alleles to obtain similar heights as the *Rht1b* mutants. An advantage of the *platz-A1* mutants is that they are already present in adapted germplasm from multiple origins, which can accelerate their deployment in breeding programs. However, the small reduction in grain size associated with the *platz-A1* mutation may constitute a disadvantage for its incorporation into commercial varieties. One strategy to compensate for this reduction is to combine the *platz-A1* mutation with the *gw-A2* mutant allele, which has been previously shown to have a positive effect on grain size in wheat (12).

In summary, this study reports the identification of *PLATZ-A1* as the causative gene of the GA-sensitive *RHT25* plant height locus and its genetic and physical interactions with DELLA. It also describes several natural loss-of-function alleles selected during wheat breeding and the molecular markers required to study their distribution and effects on grain yield in different environments. The natural and induced *PLATZ1* mutants described in this study have variable effects, which can be used to fine-tune wheat plant height in wheat breeding programs. In addition, these mutants can be combined with other GA-sensitive dwarfing genes to replace the GA-insensitive *RHT1* dwarfing genes. These new GA-sensitive semidwarf varieties will allow the exploration of the effects of deeper planting and better growth responses to nitrogen on grain yield potential in different environments.

## Materials and Methods

The mapping populations used in this study to characterize the different *RHT5* alleles are described in *SI Appendix*, Method S1. The methods for the phylogenetic analysis of PLATZ Group III proteins are described in *SI Appendix*, Method S2, whereas the EMS-induced, CRISPR-induced, and natural *PLATZ1* mutants are described in *SI Appendix*, Methods S3, S4 and S5, respectively.

The methods used for plant transformation and subcellular localization of PLATZ-A1 are described in *SI Appendix*, Method S6, whereas those used for yeast-two- and yeast-three-hybrid methods are in *SI Appendix*, Method S7. Protein Co-immunoprecipitation assays and western blotting methods are in *SI Appendix*, Method S8, and RNA extraction and real-time qRT-PCR methods are in *SI Appendix*, Method S9.

All genome coordinates and identification numbers for the annotated genes are from CS RefSeq v1.1 (37) unless indicated otherwise. Exome sequencing data was extracted from the T3/Wheat database project 2017_Wheat-CAP_UCD (https://wheat.triticeaetoolbox.org/search/genotyping_data_projects).

## Acknowledgements

The authors thank Dr. Daniel P. Woods for his help with primer design and phylogenetic analyses, Dr. Antonella Farela for providing the GRF primers, and Huiqiong Lin for her help in the Co-IP experiment. We also thank Dr. Daniel P. Woods, Huiqiong Lin, and Juan M. Debernardi for their valuable advice. We thank Nancy Blake and Dr. Luther Talbert for the Mc Neal x Thatcher mapping populations and Dr. Curtis Pozniak for providing the CDC Landmark seeds. JD acknowledges support from the USDA National Institute of Food and Agriculture the Agriculture and Food Research Initiative Competitive Grant 2022-68013-36439 (WheatCAP), and from the Howard Hughes Medical Institute.

