## Supplemental Methods and Figures for "Wheat plant height locus *RHT25* encodes a PLATZ transcription factor that interacts with DELLA (RHT1)"

**This file includes:**

Supporting Text, Methods S1 to S9

Figures S1 to S7

Supporting information References 1-20

**Other supporting materials for this manuscript include the following:**

Datasets, Tables S1 to S17 in separate Excel file

### Supporting Information Text

### Method S1. Mapping populations

For the identification of the most likely *RHT25* candidate gene, we used eight mapping populations generated from the cross between the CIMMYT line Berkut (Irene/Babax//Pastor) as the female parent and eight spring wheat lines with diverse genetic backgrounds as male parents. These lines included UC Davis varieties Patwin-515HP (PVP 2016-00390, henceforth P515HP) and Kern $_{(Yr5+Yr15+2NS+GPC\text{-}B1)}$ (subscripts indicate genes introgressed in the variety Kern, PVP 2000-00047). The non-UCD varieties included DD (Dharwar Dry), PI 70613 (LR23), RAC875, PBW343, CItr 7635 (Lr3), and RSI5$_{(Yr5+Yr15+Glu\text{-}A1a+GPC\text{-}B1)}$ (subscripts indicate genes introgressed in the variety RSI5, PI 584453).

Additional populations were generated to map and evaluate the different *PLATZ-A1* alleles. To map the loss-of-function *Rht25c* allele, we used 160 recombinant inbred lines (RILs) previously generated from the cross between common wheats McNeal and Thatcher and genotyped with SSR and DArT markers (1). The parent Thatcher has the *Rht25f* allele and McNeal the *Rht25c* allele. Lines were evaluated for plant height in the field as 3-m rows (five environments, three replications). Plant height was the average of two measurements made from the soil surface to the top of the spikes, excluding awns (1). We genotyped the population with codominant markers for *Rht25c* and the segregating *RHT-D1* gene to confirm the co-segregation of *RHT25* with one of the QTLs for plant height in this population.

To map the *Rht25d* allele identified in P515HP, we screened 75 RILs generated from the cross between Berkut (*Rht25a*) and Patwin (*Rht25d*) and identified an $F_6$ line that was still heterozygous for the *Rht25* candidate gene region but fixed for the *Rht24* region (*Rht24b*). We self-pollinated this plant to generate a Heterogeneous Inbred Family (HIF) population of 100 $F_{6:7}$ plants, which were grown in the greenhouse and evaluated for plant height.

To map the *Rht25e* allele, we developed a population by crossing RSI5 (*Rht25f*) with PI 520033 (*Rht25e*), selected an $F_4$ line fixed for *Rht24a*, and evaluated the $F_{4:5}$ in the greenhouse for plant height. We also validated the effect of the 4-bp deletion in homozygous $BC_4F_2$ lines from the cross between Chinese Spring (CS) (*Rht25e*, n= 6) and SPRING HOBBIT (*Rht25a*, n= 9).

To map the *Rht25f* allele (promoter insertion), we developed one $F_{3:4}$ and two $F_{4:5}$ populations from two different $F_2$ plants from the cross between CDC Landmark (*Rht25f*) and Berkut (*Rht25a*). All 3 populations were evaluated for plant height and peduncle length in the greenhouse.

**Method S2. Phylogenetic analysis of proteins from PLATZ Group III**

For the phylogenetic analysis, we included the 12 wheat PLATZ proteins identified in Group III in a previous study (2) and the closest rice (*Oryza sativa*), maize (*Zea mays*), and *Brachypodium distachyon* proteins. The evolutionary history was inferred using the Neighbor-Joining method (3). The percentage of replicate trees in which the associated taxa clustered together in the bootstrap test (1000 replicates) are shown next to the branches (4) in the selected optimal tree. The tree is drawn to scale, with branch lengths in the same units as those of the evolutionary distances used to infer the phylogenetic tree. The evolutionary distances were computed using the Poisson correction method and are presented in number of amino acid substitutions per site. Ambiguous positions were removed using the pairwise deletion option. We eliminated two poorly aligned regions and a total of 171 well-aligned positions were used in the final dataset (SI Appendix, Fig. S2). Evolutionary analyses were conducted in MEGA X (5).

**Method S3. *PLATZ1* EMS mutants**

To validate *PLATZ1* as the causal gene of *RHT25*, we screened the sequenced Kronos EMS mutant database (6) by BLASTN to identify loss-of-function mutations in the *PLATZ-A1* and *PLATZ-B1* homoeologs. We estimated the mutant effects using scores obtained from BLOSUM62 matrix (7) and the Sorting Intolerant from Tolerant (SIFT) (8). To obtain the SIFT scores, we blasted PLATZ1 sequence against NCBI non-redundant protein database and used the first 400 hits (the limit of the SIFT webserver) to calculate the SIFT score at the SIFT calculation server (https://sift.bii.a-star.edu.sg/www/SIFT_aligned_seqs_submit.html) (9).

Once the mutants were selected based on the two different scores, we crossed each of the mutants five times to wildtype Kronos to reduce the background mutations. We then intercrossed these plants and selected $BC_4F_1$ plants carrying both mutant alleles using molecular markers specifically designed for each mutation. We self-pollinated the double heterozygous plants and selected $BC_4F_2$ homozygous *platz-A1 platz-B1* double mutants, hereafter designated as *platz1*. We also obtained $BC_4F_2$ homozygous sister lines carrying all four possible homozygous combinations and evaluated them for plant height and peduncle length.

### Method S4. Generation of *platz-A1 platz-B1* double mutants using CRISPR-Cas9

To validate the EMS mutant results and to get truncation mutations in both *PLATZ-A1* and *PLATZ-B1*, we edited both homoeologs in the tetraploid variety Kronos using CRISPR-Cas9 (10). We designed one guide RNA targeting the third exon of both *PLATZ-A1* and *PLATZ-B1* (CTACACGATCAACAGCGCGA, *SI Appendix*, Table S2). This guide RNA was then cloned into a vector which included the Cas9 gene and a *GRF4-GIF1* chimera that increases wheat regeneration efficiency for *Agrobacterium*-mediated transformation (11). Eleven independent $T_0$ transgenic Kronos plants were obtained from the UCD Plant Transformation facility and were screened for mutations by next-generation sequencing (NGS). For the NGS screen, we used primers platz-sg2-NGS-F2 and platz-sg2-NGS-R4 (*SI Appendix*, Table S2) that can amplify both genomes. We analyzed the data using CRISgo (https://github.com/pinbo/CRISgo) and methods described previously (12). Three $T_0$ plants showed editing for both genomes and their $T_1$ plants were evaluated to confirm editing. One $T_1$ plant with confirmed editing was crossed with wildtype Kronos to segregate out the transformation vector and to create an $F_2$ population segregating for truncation mutations in both *PLATZ1* homoeologs. Finally, we characterized the effect of the different mutant combinations on plant height.

### Method S5. Characterization of *PLATZ-A1* natural variants

Using the data from exome capture, pan-genome and sanger sequencing, we found more natural variants of *PLATZ-A1* (Table 1). To test their effects on plant height, we created several populations that are described in *SI Appendix*, Table S14 and Method S1.

We used a single pair of primers (PlzAF2: CTGTATATGCATGTGTGTCTC, PlzAR1: GAGGCAGCAAAAGTGGAAGG) to genotype simultaneously the mutations present in the *Rht25b*, *Rht25c*, *Rht25d*, and *Rht25e* alleles. We run the PCR reaction at 95 °C for 3 min, followed by 38 cycles of 95 °C 20 sec, 57 °C 20 sec, 72 °C 60 sec, and one cycle at 72 °C for 7 min. We checked the PCR products in 6 % polyacrylamide gels. We used this primer pair to screen 46 *T. turgidum* ssp. *dicoccoides*, 78 *T. turgidum* ssp. *dicoccon*, and 508 durum wheat accessions from our program and from a collection described before (13) (*SI Appendix*, Table S12). For common wheat, we screened 45 lines used in the Wheat Coordinated Agricultural Project (WheatCAP), 236 photoperiod insensitive spring wheat varieties mostly from North American breeding programs (14), and 839 spring hexaploid wheat accessions from the National Small Grains wheat core collection (15) (*SI Appendix*, Table S13). We also characterized these lines for the *RHT-B1* and *RHT-D1* alleles.

### Method S6. Plant transformation and subcellular localization of PLATZ-A1

The coding sequences (CDS) of the *PLATZ-A1* gene from tetraploid wheat Kronos (*Rht25a*) without the stop codon was synthesized by GENEWIZ in a pDONR-zeo vector (hereafter, pDONR-PLATZ-A1). We transferred the pDONR-PLATZ-A1 into the binary vector pLC41 (Japan Tobacco, Tokyo, Japan) via Gateway LR reaction (https://www.thermofisher.com/us/en/home/life-science/cloning/gateway-cloning/protocols.html#lr). The resulting construct has the *PLATZ-A1* coding region driven by the maize *UBIQUITIN* (*UBI*) promoter and encodes a protein with a 3xHA tag in its carboxyl end (*UBI::PLATZ1-HA*). The UC Davis Plant Transformation Facility (http://ucdptf.ucdavis.edu/) generated five independent $T_0$ events by transforming immature embryos from Kronos with *Agrobacterium* strain EHA105 transformed with *UBI::PLATZ1-HA* using hygromycin selection. Ten $T_1$ plants for each independent event were evaluated for transgene segregation, *PLATZ1* expression levels (qRT-PCR with *ACTIN* as endogenous control), and plant height at maturity.

To test complementation of the *platz1* mutant phenotype, $T_1$ plants carrying *UBI::PLATZ1-HA* were crossed with the homozygous $BC_1F_2$ *platz1* plants. The homozygous $F_3$ plants were evaluated for plant height in a greenhouse under natural light supplemented with artificial light for 16 h, and with temperatures oscillating between 22.8 ± 1.6 °C during the day and 20.3 ± 0.3 °C during the night. We made later another transformation with a similar construct but without any tags (*UBI::PLATZ1*). We got 13 $T_0$ plants, but only 1 $T_0$ plant (202064-009) produced seeds. The other 12 $T_0$ plants were all male-sterile, and produced seed only when crossed with wildtype Kronos or Kronos homozygous $BC_4F_3$ *platz1* mutants as male parents. $T_1$ plants and $F_1$s were evaluated in the greenhouse under the same conditions as described above.

To test PLATZ-A1 subcellular localization, we transferred this gene from pDONR-PLATZ-A1 into vector pMDC84-mcherry-pUC57 by LR reaction. We transformed the p2xCa35S::mcherry-PLATZ construct into wheat protoplasts obtained from two-week-old Kronos seedlings using polyethylene glycol (PEG) as described previously (16).

**Method S7. Yeast two-hybrid (Y2H) and yeast three-hybrid (Y3H) assays between DELLA, PLATZ1 and GRF4**

For the Y2H assays, the coding sequence of *PLATZ-A1* was cloned into yeast bait vector pGBKT7 (GAL4 DNA-binding domain) between *Nde*I and *Eco*RI restriction sites using primers PLATZ-NdeI-F and PLATZ-EcoRI-R (*SI Appendix*, Table S2). The C-terminal GRAS domain of *RHT-B1* (encoding amino acids 201-621) was first cloned into pENTR/D-TOPO vector using primers RHT_Entry_F3 and RHT_Entry_R2 (*SI Appendix*, Table S2) before recombining it into yeast prey vector pLAW11 (GAL4 activation domain) by Gateway Cloning. The GRAS domain was further dissected into three sub-domains LVL (including LHRI, VHIID and LHRII domains), PFYRE and SAW. Each of these three sub-domains were cloned into pDONR/Zeo using primers listed in *SI Appendix*, Table S2 before recombining them into pLAW11 vector to generate additional prey vectors. Yeast strain Y2HGold (Clontech) was used in all yeast two-hybrid assays. The lithium acetate method was used for yeast transformation. Transformants were selected on SD medium lacking leucine (L) and tryptophan (W, abbreviated as SD-L-W) and then re-plated on SD medium lacking L, W, histidine (H) and adenine (A, abbreviated as SD-L-W-H-A) to test protein-protein interactions.

For the Y3H assays, we used the pBridge system (Clontech), which can express two proteins, a DNA-binding domain fusion, and a Bridge protein (referred as the 3rd protein). The 3rd protein is driven by an inducible promoter MET25 and can only be expressed in the absence of methionine. Addition of 1mM methionine in the medium is sufficient to inhibit its expression. The coding regions of *PLATZ-A1* and GRF4 were cloned into the pBridge vector to generate two constructs pBridge-*GRF4*-proMET25-*PLATZ* and pBridge-*PLATZ*-proMET25-*GRF4*, expressing PLATZ-A1 or GRF4 as the 3rd protein, respectively. Each of the two pBridge constructs was then paired with the AD-RhtB1-GRAS prey vector and co-transformed into yeast Gold. Transformants containing both vectors were selected on SD-L-W medium. Protein interactions were quantified using quantitative $\alpha$-galactosidase assays as described previously (17).

**Method S8. Protein Co-immunoprecipitation (Co-IP) assay and western blotting**

To validate the interaction between PLATZ-A1 and DELLA detected by Y2H, we used a Co-IP method previously described for rice (18) with minor modifications. We transformed Kronos protoplasts with 1 µg of N-terminal HA-tagged PLATZ-A1 (HA-PLATZ1) and 1 µg of C terminal cMyc-tagged DELLA (DELLA-Myc) and similar amounts of two controls, HA-PLATZ1 only, and DELLA-Myc only. We performed the transformations in a 50 ml tube containing 2 ml of protoplast (1.0 x $10^6$ per mL) each. We cultivated the transformed protoplasts in the dark at room temperature for 20 h. Total protein was extracted with 1 ml of immunoprecipitation (IP) buffer containing Thermo Scientific Halt Protease Inhibitor (Catalog number: 78425). We kept 50 µl of the raw protein extraction (for control), and used the rest for Co-IP. The pull-down assay was carried out with Thermo Scientific Pierce Anti-HA Magnetic Beads (Catalog number: 88836) overnight at 4 ºC with slow rotation. Pull-down proteins were eluted in 50 µl 1x Bio-Rad Laemmli sample buffer (Catalog number: 161-0747).

For Western Blotting, we ran 25 µl of the Co-IP elution and 5 µl of the raw protein extractions (input samples) in a 10% SDS-PAGE gel at 150 v for 1.5 h. Proteins were then transferred to a PVDF membrane using the Bio-Rad Trans-Blot Turbo Transfer System (Bio-Rad Catalog number: 1704150). The membrane was blocked with 5% non-fat dried milk dissolved in 1x TBS-T for 1 h at room temperature, and probed with an Anti-cMyc-Peroxidase monoclonal antibody (Roche 11814150001) at a dilution of 1:5000 overnight at 4 ºC. The SuperSignal West Femto Maximum Sensitivity Substrate (ThermoFisher, Catalog number 34096) was used for final detection. After imaging, the anti-cMyc antibody was removed from the membrane with the stripping buffer, re-blocked for 1 h at room temperature, and then probed with an HRP-conjugated HA Epitope Tag monoclonal antibody (Invitrogen, 26183-HRP) at a dilution of 1:5000 for 1 h at room temperature.

**Method S9. RNA extraction and real-time qRT-PCR analysis**

RNAs from various tissues were extracted using the Spectrum Plant Total RNA Kit (Sigma-Aldrich, STRN50). One µg of RNA was treated with RQ1 RNase-Free DNase (Promega, M6101) first and then used for cDNA synthesis with the High-Capacity cDNA Reverse Transcription Kit (Thermo Fisher Scientific, 4368814). The cDNA was then diluted 10 times and 4 µl of the dilution was used as template in the qRT-PCR assays. We performed the quantitative reverse transcription-PCR (qRT-PCR) assays using Quantinova SYBR Green PCR kit (Qiagen, 208052) in a 7500 Fast Real-Time PCR system (Applied Biosystems). For PCR reactions, we used one cycle at 95 °C for 2 min and 40 cycles of 95 °C for 5 s and 60 °C for 30 s, followed by a melting curve program. We quantified expression using the delta Ct method ($2^{-\Delta CT}$) with *ACTIN* as endogenous control (19). Primers for *PLATZ1* amplify both the A and B genome homoeologs and have an efficiency of 98% (*SI Appendix*, Table S2).

**Supporting Figures**

**Fig. S1**. Phylogenetic analysis of PLATZ proteins from Group III. The study includes 12 wheat PLATZ proteins from Group III together with their closest rice (Os), maize (Zm), Brachypodium (BRADI), *Pisum* (PsPLATZ1), and Arabidopsis (AtORESARA15) proteins. The evolutionary history was inferred using the Neighbor-Joining method as described in *SI Appendix*, Method S2. We eliminated two poorly aligned regions and a total of 171 well-aligned positions were used in the final dataset (*SI Appendix*, Fig. S2). Names of the wheat genes are based on *SI Appendix*, Table S3.

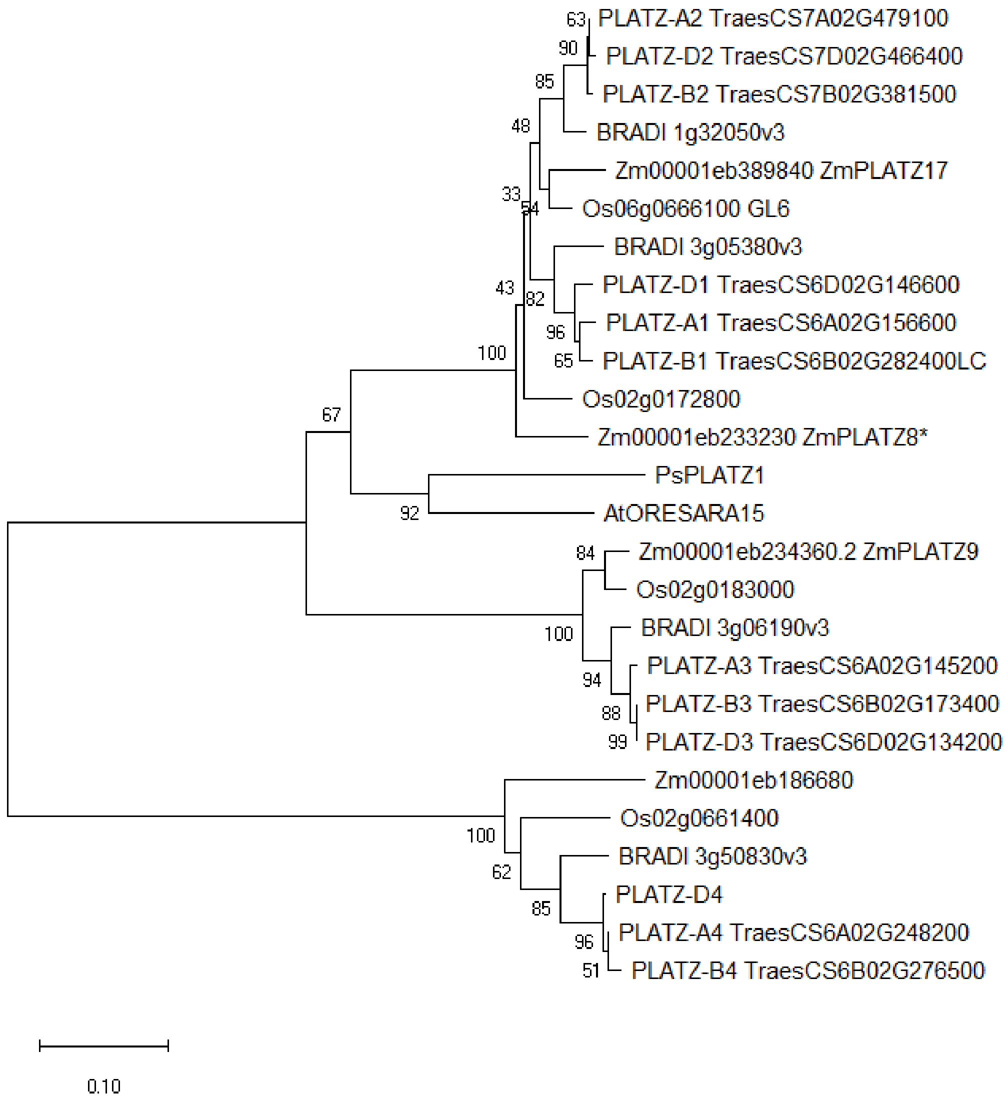

**Fig. S2**. Alignment of twenty-six PLATZ proteins from Group III, including 12 wheat proteins from Group III (2) and the closest rice (Os), maize (Zm), *Brachypodium* (BRADI), *Pisum* (Ps) and Arabidopsis (At) proteins. Poorly aligned regions were eliminated in regions indicated by ^ below the sequence. Sequences were aligned using MUSCLE algorithm in MEGA X. The two non-canonical zinc fingers are indicated in red below the sequences.

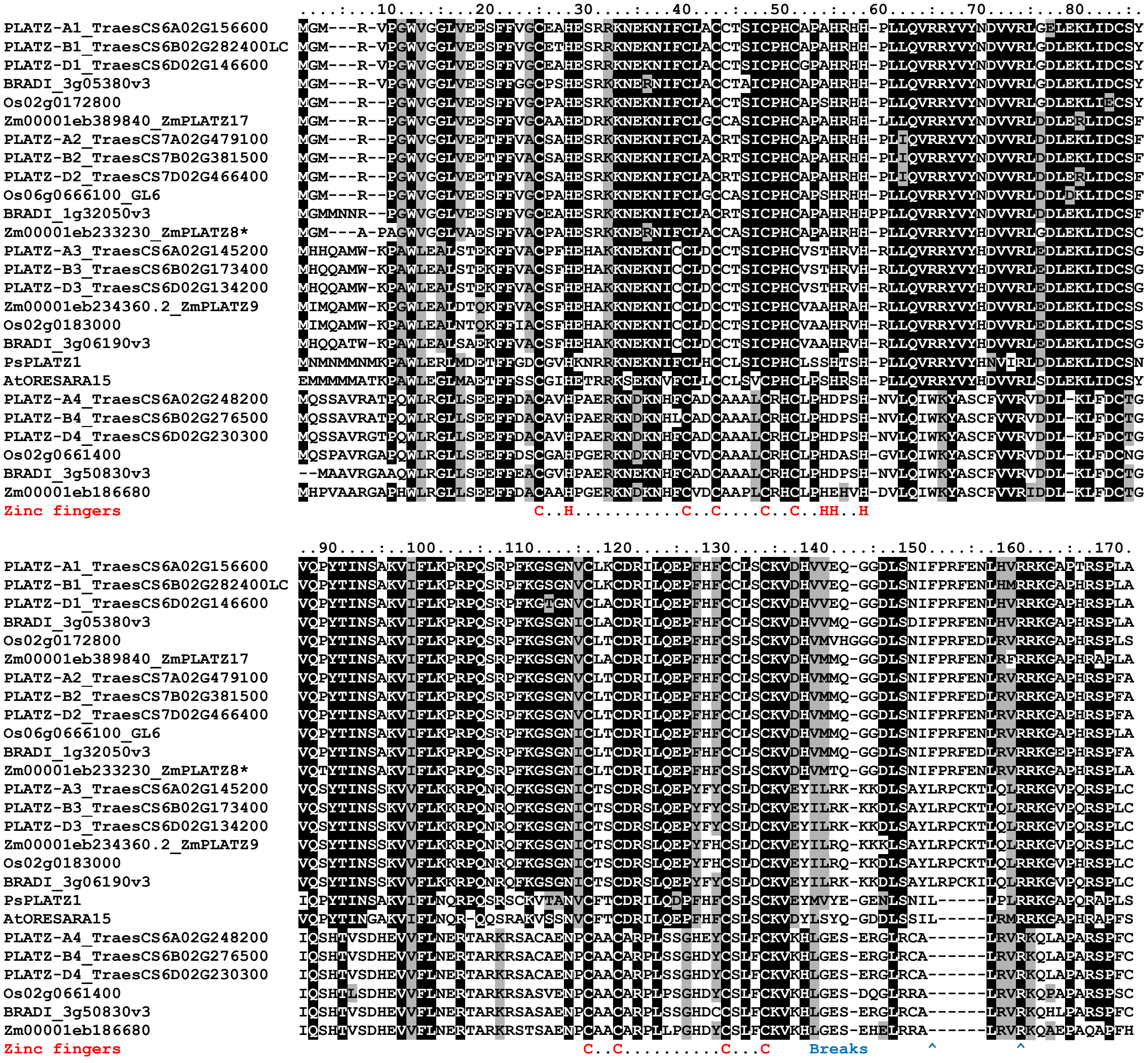


The following sequences are incorrectly annotated in the current genomes and were modified as follows:

- *PLATZ-B1*: in Chinese Spring the last 10 bp are missing (region with multiple Ns) so we replaced them by the corresponding sequence in Kronos and other wheat genomes.
- *ZmPLATZ8*: we added a missing 1st exon based on more recent annotation *Zm00001eb233230.*
- *Zm00001eb186680*: protein was truncated due to an incorrectly annotated splicing site at the end of exon 3. We moved this splicing site 14 bp earlier and restored the reding frame of exon 4.
- Os02g0661400: we eliminated 51 bp incorrectly annotated at the start of exon 3, which would have translated into 17 amino acids that are not present in any other PLATZ protein from Group III.

**Fig. S3.** Induced mutations in *PLATZ1*. (**A**) Ethyl methane sulfonate (EMS) induced mutations in tetraploid wheat Kronos are indicated below the *PLATZ-A1* and *PLATZ-B1* genes with blue arrows. Induced deletions and insertions generated by CRISPR-CAS9 are indicated by red arrows on top of the genes. Boxes with numbers indicate the four exons, characteristic of all *PLATZ* genes from Group III. (**B**) Effect of the mutation in the acceptor splice site of the third intron of *PLATZ-A1* in Kronos mutant line K2557 on RNA splicing products. RT-PCR of *PLATZ-A1* using RNA samples extracted from leaves of wildtype Kronos resulted in a 339 bp band (expected from correct splicing), whereas those extracted from mutant line K2557 showed a strong 467 bp band (generated by intron 3 retention) and a fainter 328 bp band (generated by a 21 bp deletion from exon 4 as a result of the utilization of an alternative splicing acceptor site).

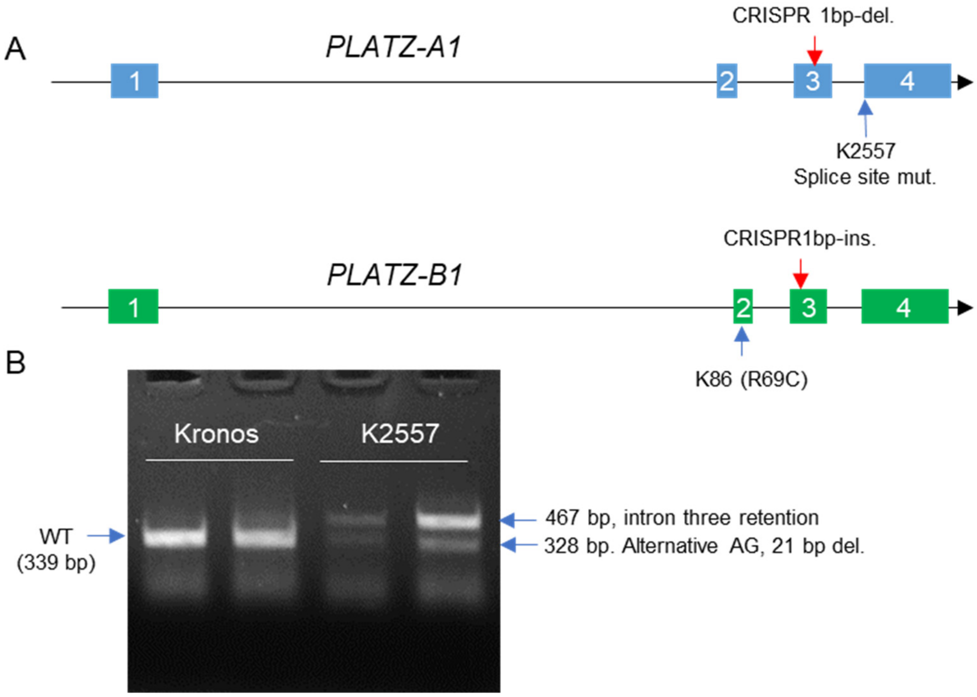

**Fig. S4.** Effect of CRISPR-CAS9 induced indels in *PLATZ1*. (**A**) Differences in plant height in selected sister $F_3$ plants homozygous for the wildtype (WT) and *CRplatz1* indels in both *PLATZ1* homoeologs. (**B**) Interaction graph for plant height for the four possible homozygous classes combining the *CRplatz-A1* and *CRplatz-B1* indels. (**C**) Interaction graph for peduncle length. ns = not significant, * = $P < 0.05$, and ** = $P < 0.01$.

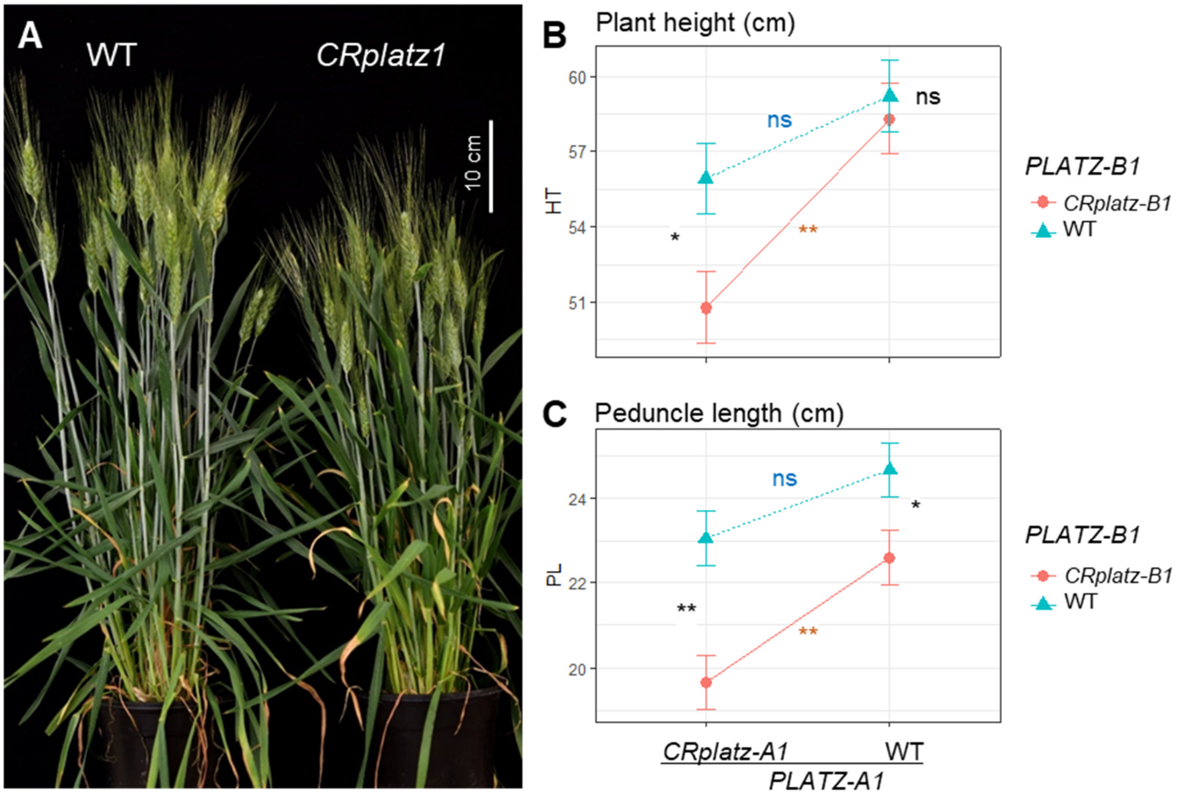

**Fig. S5.** Expression of *PLATZ1* in UBI::PLATZ-A1:HA transgenic plants (Y) and non-transgenic sister controls (N). RNA samples were extracted from leaves of 16-day-old individual $T_1$ plants segregating for the transgene in five independent events. Primers amplify both the transgene and the endogenous *PLATZ-A1* and *PLATZ-B1* genes (*SI Appendix*, Table S2). Transcript levels were calculated using the delta Ct method using *ACTIN* as endogenous control (*SI Appendix*, Method S9).

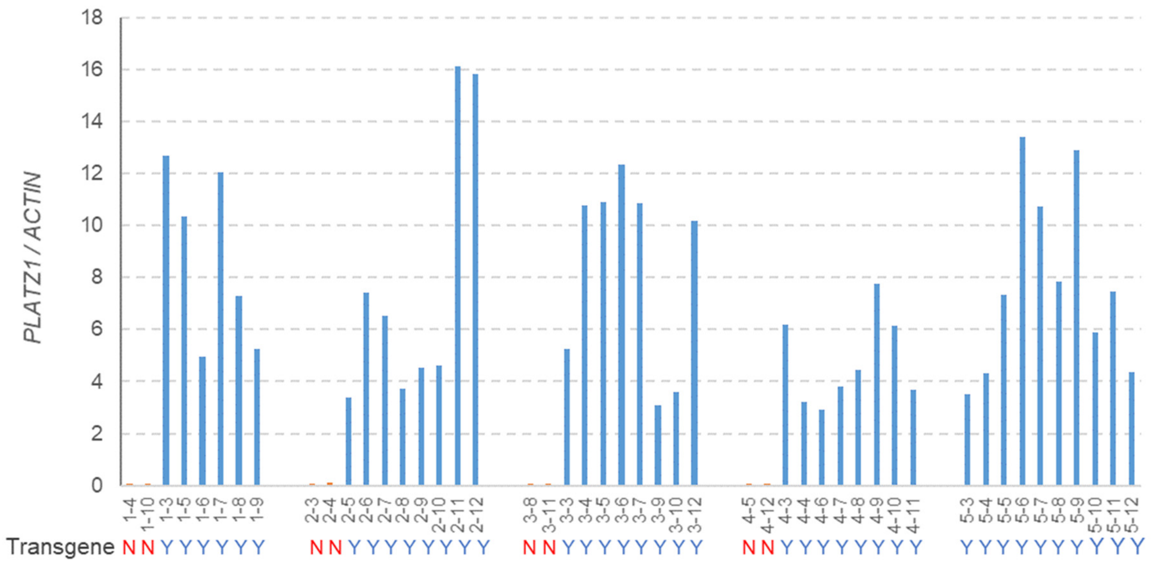

**Fig. S6.** Relative expression of *PLATZ1* homoeologs in **A**) hexaploid wheat Chinese Spring, RNA-seq transcripts per million (TPM) in different tissues and developmental stages (20), and **B**) tetraploid wheat Kronos RNA-seq TPM in spike development time course (21). Only uniquely mapped transcripts are presented (22). "Z" numbers represent developmental stages in Zadoks' scale (23). **C**) Transcript levels of *PLATZ1* (both homoeologs) in different sections of the elongating peduncle at ear emergence (Zadok 55): node, 1 cm above node (Int. 1cm), and 2 cm above the previous region (Int. 2cm, n = 4, with each replication representing a pool of three peduncles). qRT-PCR using the Delta Ct method to calculate transcript levels of *PLATZ-A1* relative to the *ACTIN* endogenous control. Different letters over the bars indicate significant differences in Tukey test ($P < 0.05$). The raw data and statistics are available in *SI Appendix*, Table S8.

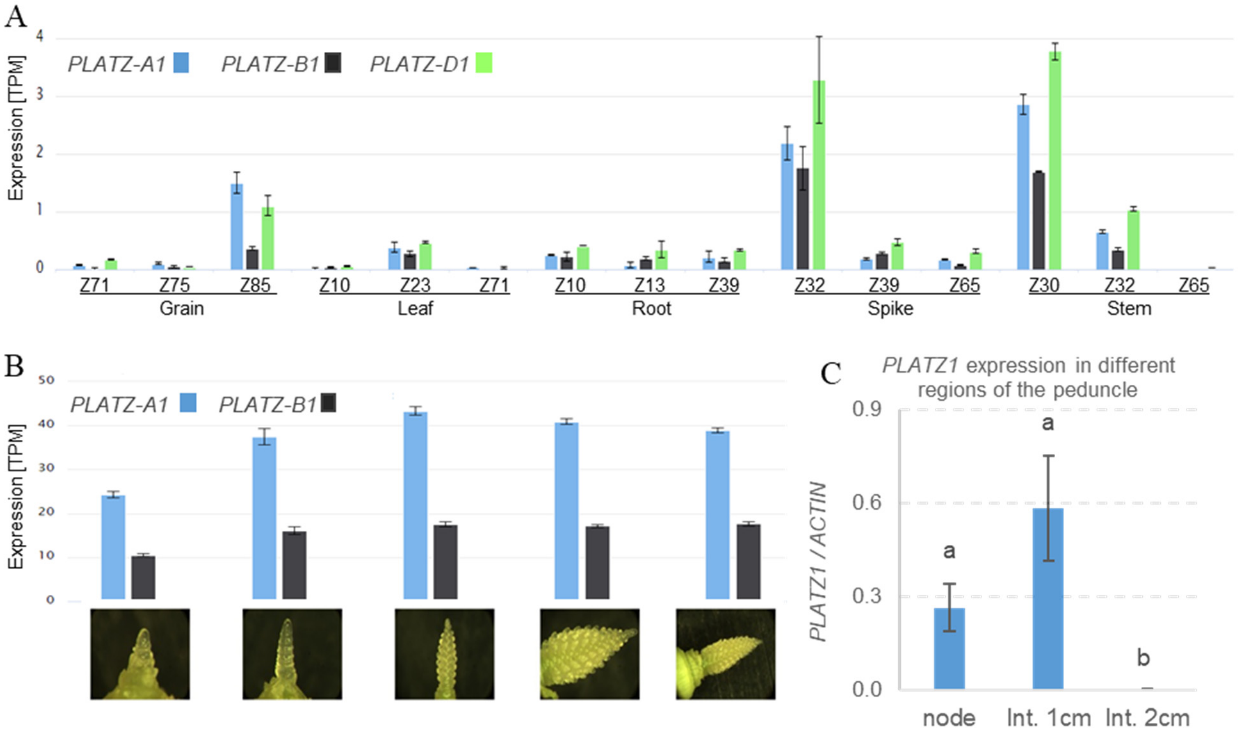

**Fig. S7.** Natural variation in the coding region of *PLATZ-A1*. *PLATZ-A1* natural variants detected using primer pairs. **A-B**) PlaAF2/PlzAR1 (*Rht25a* to *e*) and **C**) platz6A-CDC-F1/R1 (*Rht25f*) (*SI Appendix*, Table S2). **A-B**) Berkut has the *Rht25a* allele (WT, 154 bp PCR product), Chinese Spring the *Rht25e* allele (4-bp deletion, 150 bp PCR product), UC1110 the *Rht25b* allele (13-bp deletion, 141 bp PCR product), McNeal the *Rht25c* allele (19-bp deletion, 135 bp PCR product), P515HP the *Rht25d* allele (splice site mutation, 154 bp PCR product, digested in 21 +133 bp fragments by *Sau*96I digestion). **B**) Digestion with *Sau*96I helps differentiate the splice site mutation in *Rht25d*. **C**) Berkut (*Rht25a*) x CDC Landmark (*Rht25f*) $F_2$ plants. The lower band corresponds to the *Rht25a* allele (157 bp) and the upper band to the *Rht25f* allele (541 bp due to a 384-bp insertion in the *PLATZ-A1* promoter).

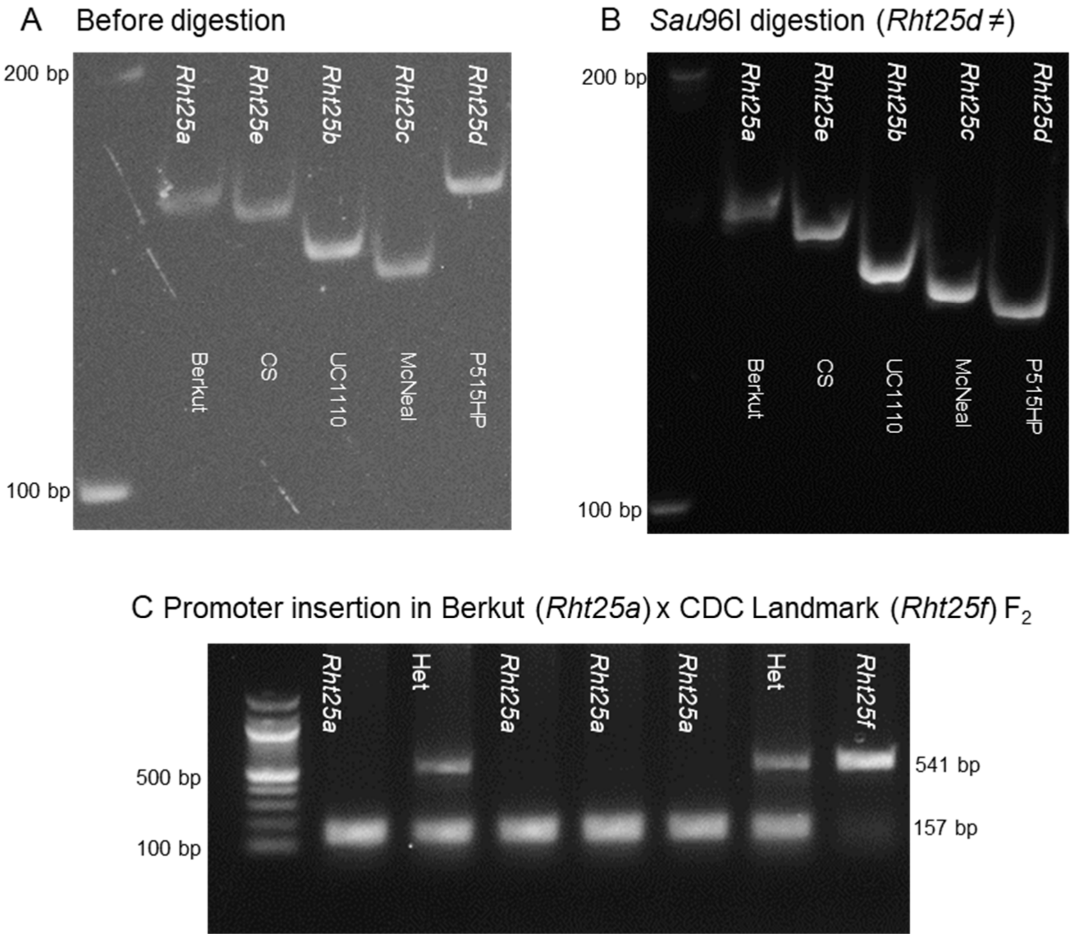

**Supporting information references**


1. J. D. Sherman, J. M. Martin, N. K. Blake, S. P. Lanning, L. E. Talbert, Genetic basis of agronomic differences between a modern and a historical spring wheat cultivar. *Crop Sci.* **54**, 1-13 (2014).
2. Y. X. Fu *et al.*, Identification and characterization of PLATZ transcription factors in wheat. *Int. J. Mol. Sci.* **21**, 8934 (2020).
3. N. Saitou, M. Nei, The Neighbor-Joining method - a new method for reconstructing phylogenetic trees. *Mol. Biol. Evol.* **4**, 406-425 (1987).
4. J. Felsenstein, Confidence-limits on phylogenies - an approach using the bootstrap. *Evolution* **39**, 783-791 (1985).
5. S. Kumar, G. Stecher, M. Li, C. Knyaz, K. Tamura, MEGA X: Molecular evolutionary genetics analysis across computing platforms. *Mol. Biol. Evol.* **35**, 1547-1549 (2018).
6. K. V. Krasileva *et al.*, Uncovering hidden variation in polyploid wheat. *Proc Natl Acad Sci U S A* **114**, E913-E921 (2017).
7. S. Henikoff, J. G. Henikoff, Amino-acid substitution matrices from protein blocks. *Proc Natl Acad Sci U S A* **89**, 10915-10919 (1992).
8. P. C. Ng, S. Henikoff, Predicting deleterious amino acid substitutions. *Genome Res.* **11**, 863-874 (2001).
9. N. L. Sim *et al.*, SIFT web server: predicting effects of amino acid substitutions on proteins. *Nucleic Acids Res* **40**, W452-457 (2012).
10. M. Jinek *et al.*, A programmable dual-RNA-guided DNA endonuclease in adaptive bacterial immunity. *Science* **337**, 816-821 (2012).
11. J. M. Debernardi *et al.*, A GRF-GIF chimeric protein improves the regeneration efficiency of transgenic plants. *Nat. Biotechnol.* **38**, 1274-1279 (2020).
12. J. Zhang, *Check CRISPR editing events in transgenic wheat with next-generation sequencing*. S. H. Wani, A. Kumar, Eds., Springer Protocols Handbooks. Genomics of Cereal Crops (Springer US, New York, NY, 2022).
13. S. Chao *et al.*, Evaluation of genetic diversity and host resistance to stem rust in USDA NSGC durum wheat accessions. *Plant Genome* **10** (2017).
14. J. L. Zhang *et al.*, Identification and validation of QTL for grain yield and plant water status under contrasting water treatments in fall-sown spring wheats. *Theor. Appl. Genet.* **131**, 1741-1759 (2018).
15. M. Maccaferri *et al.*, A genome-wide association study of resistance to stripe rust (*Puccinia striiformis* f. sp. *tritici*) in a worldwide collection of hexaploid spring wheat (*Triticum aestivum* L.). *G3 (Bethesda)* **5**, 449-465 (2015).
